# Norbornene homopolymerization limits cell spreading in thiol-ene photoclick hydrogels

**DOI:** 10.1101/2025.04.18.648735

**Authors:** James L. Gentry, Steven R. Caliari

## Abstract

Thiol-ene click chemistry is a powerful tool for designing hydrogels mimicking the mechanical and biochemical properties of 3D cellular microenvironments. The high selectivity of thiol-norbornene step-growth polymerization enables precise control of crosslinking mechanism, circumventing the alkene homopolymerization present in other systems that can prevent encapsulated cell spreading. Limited stress relaxation of a dynamically-crosslinked norbornene-modified hyaluronic acid (NorHA) hydrogel employing a thiol-norbornene photoclick reaction led us to investigate the prevalence of norbornene homopolymerization in this supposed click reaction. Norbornene conversion was quantified in multiple thiol plus norbornene-modified polymer system permutations, revealing higher norbornene conversion than expected for 1:1 thiol-ene addition. We showed that decreasing the number of norbornenes per NorHA chain (f) mitigated network formation via norbornene homopolymerization. Dynamic hydrogels fabricated with NorHA of f = 8 (Nor_8_HA) exhibited 93.0 ± 1.6% relaxation, while those fabricated with NorHA of f = 40 (Nor_40_HA) achieved only 42.3 ± 0.1% relaxation. As early as day 3 of culture, Nor_8_HA hydrogels facilitated spreading of encapsulated human mesenchymal stromal cells (hMSCs) into a spindle-like morphology (aspect ratio: 2.95 ± 0.38), while Nor_40_HA hydrogels appeared to constrain cells into a spherical or compact star morphology (aspect ratio: 1.22 ± 0.01). Inference of a single-cell morphological space derived from a shape-matching distance metric validated the two distinct hMSC morphological phenotypes primarily associated with polymer f. Despite its widespread use as a click reaction, radical-mediated thiol-norbornene crosslinking was found to not be stoichiometric in dilute aqueous conditions used to fabricate hydrogels. Altering network topology through polymer f enabled the rescue of hydrogel dynamic behavior and encapsulated hMSC spreading, despite the presence of norbornene homopolymerization, highlighting the need to consider network-level properties when designing engineered cellular microenvironments.

## INTRODUCTION

Click chemistry has proven to be an invaluable design tool for introducing diverse functions into hydrogel matrices^1^. Click reactions are characterized by their high efficiency and specificity, often achieving near-stoichiometric conversion^2^. They have been used to tailor drug release kinetics out of hydrogel depots^3^, modify hydrogel surface properties^4,5^, and introduce shape memory behavior for application as stimulus-responsive hydrogel actuators^6^. Many of these reactions are non-cytotoxic and bio-orthogonal, making them particularly advantageous for cell encapsulation within hydrogels. Hydrogels have since emerged as useful substrates for modeling cellular microenvironments *in vitro*, offering researchers the ability to recapitulate key aspects of native tissues like mechanics, bioactivity, and remodeling capability^7^.

Tissues enable vital cell processes like spreading and migration by providing attachment points to move through 3D space and by allowing substrate remodeling through degradation or physical deformation. The development of hydrogels as synthetic 3D cellular microenvironments has enabled investigations of the role of tissue properties, like mechanics, on cell behavior without confounding factors present *in vivo*. Unfortunately, conventional hydrogel designs that include static covalent crosslinks create restrictive environments that limit cell spreading and other normal cell behaviors^8^. To overcome this limitation, hydrogels permissive to cell spreading can be engineered through the inclusion of dynamic non-covalent interactions to mimic the viscoelasticity of natural tissue^9^ and/or the inclusion of proteolytically-degradable crosslinkers to enable cell-mediated hydrogel remodeling^10^. Permissive 3D cellular micro-environments have been demonstrated to dictate stem cell lineage commitment^8,9,11^, facilitate patterning and morphogenesis in developmental models^12,13^, and encourage tissue regeneration^10,14^. Substrate permissivity is also important when modeling disease states where tissue mechanics change and directly influence cell behavior, such as fibrosis^15–17^ and cancer^18^. Together, these studies highlight the importance of designing permissive hydrogels for diverse biomedical applications.

When developing permissive hydrogels, formation of static non-degradable covalent bonds must be minimized. Click reactions are an obvious choice to precisely control crosslinking mechanism. Among these, the thiol-norbornene photoclick reaction has gained prominence due to its ability to afford spatiotemporal control of crosslinking via light as well as its relative insensitivity to oxygen, fast kinetics, and high thiol-ene selectivity with limited norbornene homopolymerization^19–21^. This reaction scheme has garnered widespread use in the design of permissive hydrogels that are proteolytically-degradable^13,20,22–31^ or viscoelastic^32,33^. Thiol-norbornene crosslinking has also been used to create shape-memory^34,35^ and mechanically dynamic^36^ substrates. Additionally, thiol-norbornene click reactions can be used to introduce bioactive molecules to the network, not just crosslinks. Bioactive molecule presentation can be spatiotemporally controlled with light as demonstrated with adhesive ligands and growth factor mimetics^37,38^.

Successful fabrication of such complex photopatterned hydrogels is more easily achieved when using a polymer with high degree of functionalization (f). At the same concentration of functional groups, high f polymers require fewer crosslinks for network formation to occur compared to low f polymers like 4-arm PEG^21,39^. This allows a greater proportion of norbornenes to be consumed by non-cross-linking thiols while maintaining gelation capability. However, more efficient network formation through intended crosslinks also means that unintended off-target crosslinks can easily induce gelation as well. While radical-mediated thiol-norbornene addition is classified as a click reaction, low levels of norbornene homopolymerization have been demonstrated, especially in the absence of thiols^19^. Static crosslinks created via norbornene homopolymerization could limit a network’s dynamic behavior achieved through degradability and/or viscoelasticity, inhibiting cell spreading and preventing the use of these hydrogels as engineered 3D cellular microenvironments.

To determine if norbornene homopolymerization could limit network remodeling of a dynamic hydrogel, we utilized a norbornene-modified hyaluronic acid (NorHA) hydrogel that employs a thiol-norbornene photoclick reaction to create physical crosslinks. We validated the presence of norbornene homopolymerization and demonstrated its effects on network stress relaxation by modulating polymer f. Morphology of encapsulated cells is similarly shown to be dependent on polymer f.

## RESULTS AND DISCUSSION

### A photoclick version of a supramolecular guest-host hydrogel unexpectedly exhibits incomplete stress relaxation

We designed a stress relaxing hydrogel system leveraging supramolecular adamantane-β-cyclodextrin guest-host chemistry^40^ to evaluate how norbornene homopolymerization might impact viscoelasticity. In the conventional version of this system, mixing guest adamantane-modified HA (AdHA) and host β-cyclodextrin-modified HA (CDHA) leads to the spontaneous formation of a stress relaxing hydrogel whose rheological properties are governed by the reversible physical interactions between adamantane and β-cyclodextrin (**Fig. 1a**, left). We would expect this hydrogel to exhibit nearly complete stress relaxation with a loss factor (loss modulus (G”)/storage modulus (G’)) approaching 1. Our lab has previously developed a photocurable version of this system that utilizes CDHA, NorHA, and a thiolated adamantane peptide (**Fig. 1a**, right)^33^. This system was designed to exhibit the same stress-relaxing properties of the spontaneously gelling CDHA & AdHA system but with more homogenous mechanical properties and improved tunability, as the guest-host complex does not act as a cross-link prior to the photoclick thiol-ene reaction. If norbornene homopolymerization significantly contributes to network formation through unintended covalent crosslinking, then the photocurable system should exhibit reduced extent of stress relaxation and loss factor compared to the spontaneously gelling system.

**Figure 1.**
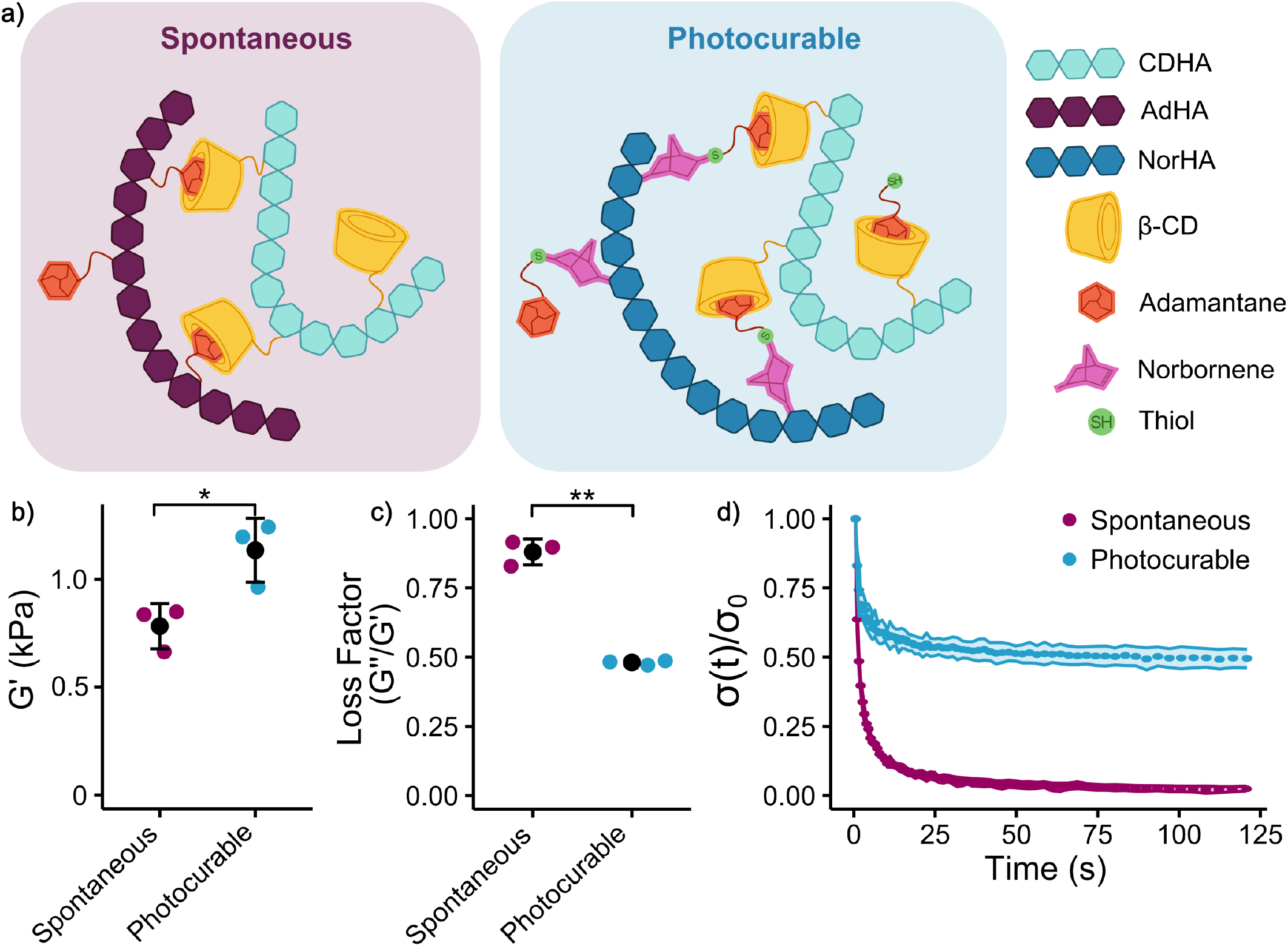
Photocurable supramolecular hydrogels show reduced stress relaxation compared to spontaneous supramolecular hydrogels. a) Schematics showing the crosslinking mechanism of a spontaneously gelling hydrogel (left) and a radical-initiated photoclick hydrogel (right). For the spontaneous system, two macromers functionalized with either β-cyclodextrin (CDHA) or adamantane (AdHA) are mixed, with the complexes between the two groups acting as crosslinks. For the photocurable version, CDHA is pre-complexed with thiolated adamantane. This macromer is then mixed with a norbornene-functionalized macromer (NorHA) where radical-initiated thiol-ene addition creates the physical crosslinks similar to the spontaneous system. Spontaneous and photocurable hydrogels with the same concentrations of adamantane and βcyclodextrin were fabricated. While the mechanics of the two systems were expected to be similar, the b) storage modulus (G’) of the photocurable version was significantly higher than the spontaneous version, and the c) loss factor (G”/G’) was significantly reduced. d) Similarly, the photocurable version exhibited only partial stress relaxation (stress over time, σ(t), normalized to initial stress, σ(0)) while the spontaneous version completely relaxed as expected. *N* = 3 hydrogels per group. Statistical analyses performed via Welch’s t-tests. **: *p* < 0.01, *: *p* < 0.05.

Viscoelastic behavior of both systems was determined via shear oscillatory rheology. The spontaneously gelling hydrogels were fabricated prior to rheological characterization by mixing CDHA (22% modified, see **Fig. S1**) and AdHA (26% modified, see **Fig. S2**) solutions at concentrations giving a 1:1 molar ratio of adamantane:β-cyclodextrin. This formulation was mimicked in the photocurable version by maintaining the CDHA concentration and molar concentration of adamantane via the thiolated adamantane. NorHA (24% modified, see **Fig. S3**) was added at a concentration such that there was a 1:1 molar ratio between thiolated adamantane (see chemical structure at **Fig. S4**) and norbornene. These hydrogels were fabricated *in situ* on the rheometer.

Despite having less polymer and theoretically larger mesh size, the photocurable system had a storage modulus (G’) almost 1.5 times that of the spontaneous system (*p* = 3.25×10^-2^, **Fig. 1b**). The spontaneously gelling system had a loss factor close to 1 as previously reported (0.880 ± 0.046)^40^, while the photocurable system exhibited a significantly lower loss factor of 0.481 ± 0.008, suggesting the presence of stable covalent crosslinks (*p* = 3.65×10^-3^, **Fig. 1c**). Stress relaxation behavior further supported this hypothesis as the spontaneously gelling system fully relaxed (98% relaxation) and the photocurable system could only achieve 51% relaxation, suggesting that stress is stored by crosslinks with greater stability than guest-host complexing (*p* = 8.79×10^-4^, **Fig. 1d**).

To verify that crosslinks other than the guest-host complexes were present in the photocurable system, these hydrogels were incubated overnight in excess adamantane to disrupt the supramolecular crosslinks. The spontaneously gelling hydrogels lost structural integrity and coated the bottom of the incubation dish (**Fig. S5**). However, the photocured hydrogels maintained their shape, demonstrating the presence of some other crosslinking mechanism.

### Norbornene conversion greatly exceeds thiol concentration, suggesting non-trivial rates of norbornene homopolymerization

To determine if norbornene homopolymerization could be driving covalent network formation, we exposed a solution containing only NorHA and lithium acylphosphinate (LAP) photoinitiator to ultraviolet (UV) light for 5 minutes, after which a robust hydrogel had formed. NMR spectroscopy comparing unreacted NorHA to this sample showed a 77% reduction in norbornene peak area (**Fig. S6**), suggesting norbornene homopolymerization as the crosslinking mechanism. Increased LAP concentration demonstrated that hydrogels upwards of G’ ∼ 1.5 kPa could be fabricated without dithiol crosslinkers (**Fig. 2a**).

**Figure 2.**
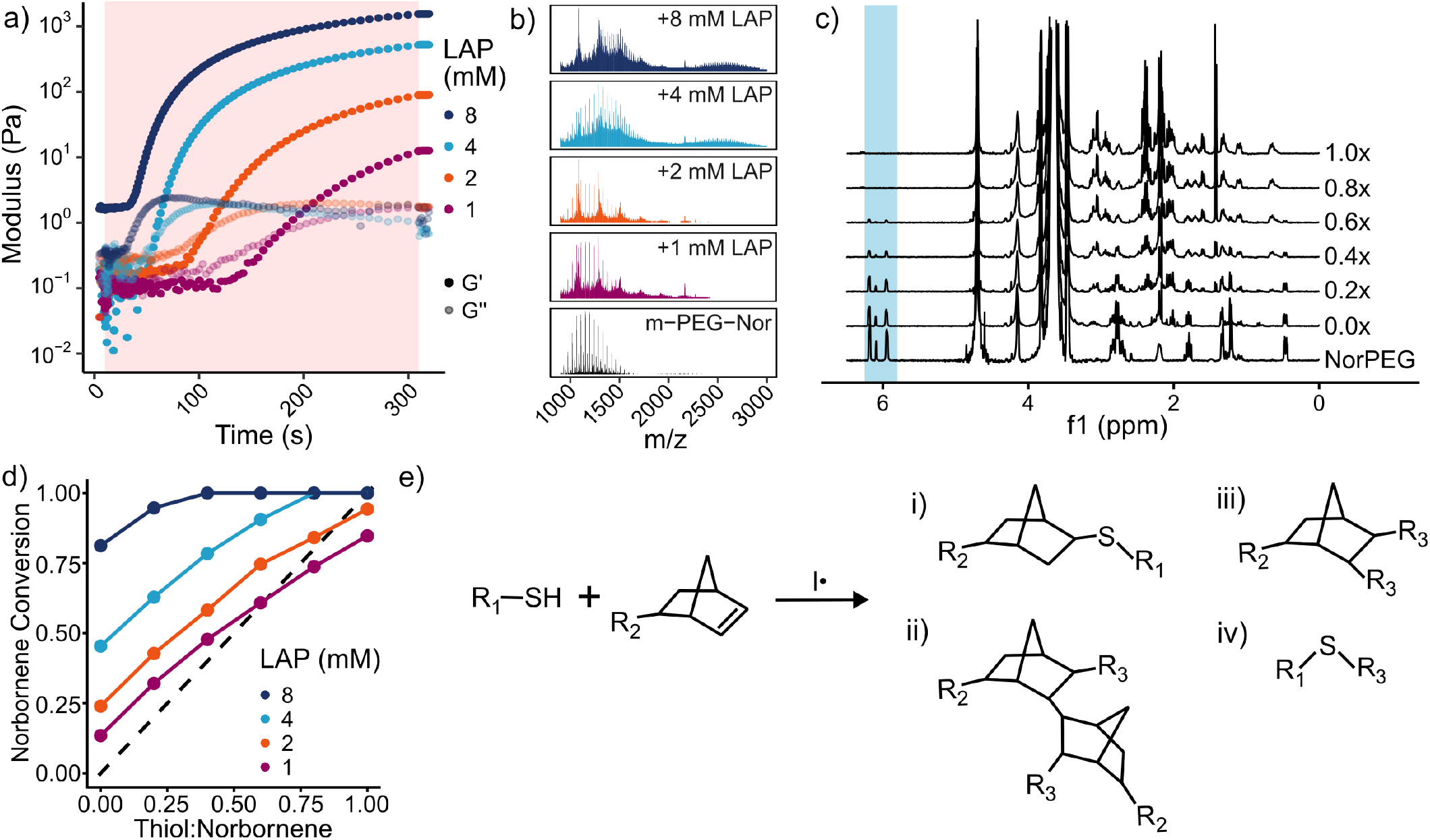
Evaluating thiol-norbornene addition stoichiometry in dilute aqueous conditions. a) UV curing of NorHA in the presence of LAP photoinitiator induced gelation at all tested LAP concentrations, suggesting that norbornene homopolymerization is prevalent enough to drive network formation. Storage modulus reached upwards of 1.5 kPa in the 8 mM LAP condition. The red rectangle indicates UV exposure time. b) MALDI-ToF spectra of 11 mM 1 kDa mono-substituted PEG-norbornene (m-PEG-Nor) before (bottom) and after exposure to UV light in the presence of LAP photoinitiator. Norbornene homodimers appear at high LAP to norbornene ratios. c) ^1^H NMR spectra of NorPEG (11 mM norbornene) reacted with increasing amounts of cysteine in the presence of 4 mM LAP photoinitiator. Reduction in norbornene peak area (δ = 5.80-6.25), even with no thiol present (0.0x), is apparent when compared to unreacted NorPEG (bottom spectrum). d) Norbornene conversion as a function of thiol:norbornene ratio for different LAP concentrations is plotted as actual conversion (change in NMR peak area) against expected conversion (thiol concentration). Norbornene conversion generally exceeded the expected amount. Increased initiator to norbornene ratio resulted in even higher excess conversion, indicating that norbornene conversion partially occurs via a route other than thiol-ene addition. e) Revised thiol-norbornene polymerization scheme in dilute aqueous conditions at higher initiator concentrations used for hydrogel fabrication. Products include non-trivial amounts of i) the desired thiol-ene addition, ii) norbornene homopolymer, iii) radical terminated norbornene, and iv) radical terminated thiol. R_3_ indicates hydrogen or any radical forming species in the solution, including initiator.

To more concretely determine the presence of norbornene homopolymerization and the degree of polymerization, we performed MALDI-ToF on a mono-substituted methyl-PEG-norbornene polymer after exposing to UV light in the presence of LAP photoinitiator (**Fig. 2b**). After UV exposure, the spectra were less regularly spaced compared to the unreacted PEG, indicating the presence of a wide variety of products. As LAP concentration increased, the PEG’s mass shifted right, suggesting radical combination with the norbornene. At 4 and 8 mM LAP, PEG dimerization was apparent. We did not find evidence of homopolymer chain length greater than 2 even after increasing laser power (data not shown).

The above experiments validated that norbornene homopolymerization could cause network formation in the extreme case of absence of thiol. Since thiol-ene addition is reportedly much more favorable than norbornene homopolymerization^19^, we next investigated if excess norbornene conversion also occurred in the presence of thiols. Solutions containing 4-arm norbornene-modified polyethylene glycol (NorPEG) at 11 mM norbornene, 4 mM LAP, the reducing agent TCEP, and increasing amounts of thiol in the form of cysteine were UV cured and prepared for ^1^H NMR. Thiol concentration was set relative to norbornene concentration (i.e., 0.6x indicates a 0.6:1 molar ratio of thiol:norbornene). Thioether formation at δ = 2.4-2.6 ppm in a similar NorPEG hydrogel system was previously used to validate the efficiency of the thiol-norbornene addition^20^.

As expected, norbornene peak area (δ = 5.80-6.25 ppm) decreased even in the absence of thiol, and it continued to decrease as thiol concentration increased (**Fig. 2c**). Norbornene conversion exceeded the expected 1:1 thiol:norbornene conversion of a thiol-ene click reaction at every thiol concentration, indicating high concentrations of either norbornene homopolymers or radically terminated norbornene groups (**Fig. 2d**). Thioether peaks were not apparent, so the relative contributions of thiol-ene or ene-ene additions to norbornene conversion were not determined. Norbornene conversion did show photoinitiator concentration-dependent behavior, with more photoinitiator resulting in higher conversion. While lowering initiator concentration likely reduces norbornene homopolymerization, it also results in incomplete norbornene conversion at 1.0x thiol concentration. Low initiator concentration reduces termination rate, but also reduces reaction rate and therefore increases gelation time^41^. Perhaps sub mM LAP concentration and long UV exposure times could still yield a robust hydrogel with click-like reaction efficiency. However, such an approach would prevent quick gelation and expose cells to harmful light irradiation for extended times, nullifying advantages that make this photocurable system attractive for bioengineering applications such as 3D cell culture and bioprinting^42^.

To assess reproducibility of this behavior across polymer systems and thiol species, we repeated this experiment with NorHA and either cysteine or thiolated adamantane (**Fig. S7**). Since the thiolated species were monosubstituted, no crosslinks should form during the thiol-ene reaction. Nevertheless, all samples regardless of thiol concentration gelled, necessitating a hyaluronidase digestion step prior to ^1^H NMR analysis. The additional systems exhibited similar amounts of excess norbornene conversion (**Fig. S8**). Within system reproducibility was assessed by repeating the NorHA experiments, which showed near-identical conversions (**Fig. S9**). The complexity of the NorHA and thiolated adamantane spectra again prevented identification of thioether peaks.

We attempted to monitor the thiol and norbornene conversion in real time with FTIR as done in concentrated pure polymer solutions^19,43^. The water in the samples overwhelmed the signals of interest (thiol at 2570 cm^-1^ and alkene of norbornene at 713 cm-^1^). Even after solvent blanking, peaks at these positions were not apparent. Increasing polymer and thiol concentration would allow for improved signal but change the dilute aqueous conditions relevant to hydrogel formation that we are investigating.

In contrast to other studies^20,21^, we found that radical-initiated thiol-norbornene addition does not fulfill the high efficiency and minimal byproducts criteria of a click reaction in dilute aqueous conditions. The thiol-norbornene click reaction was originally described in a solventless system using pure liquid polymers at molar concentrations^19^. The ratio of initiator to norbornene was limited to roughly 1:1000 to prevent termination by radical addition, in contrast to the roughly 1:10 ratio typical in hydrogel systems. Premature termination was prevented at these low initiator concentrations due to the absence of any other radical-forming species in the system. However, the abundance of radical forming species in aqueous solution necessitates the use of high initiator concentrations to combat loss of initiator efficiency. Conversion of norbornene and thiol is therefore in part due to homopolymerization and/or termination via radical combination, not solely thiol-norbornene step-growth polymerization (**Fig. 2e**).

### Reduction in functional groups per polymer (f) enables complete stress relaxation of the photoclick supramolecular hydrogel

With the contribution of norbornene homopolymerization to network formation confirmed, we wanted to identify a way to mitigate its effects on hydrogel properties. The rate of norbornene homopolymerization could be decreased by increasing the concentration of thiols, thereby promoting thiol-norbornene addition. Oversaturating the norbornenes with 3x excess thiols reduced covalent network formation (**Fig. S10**), but this approach was wasteful and did not allow for easy incorporation of other thiolated molecules.

We instead turned to altering network-level properties to circumvent network formation via norbornene homopolymerization. Changing the number of functional groups on a polymer (f) significantly affects network topology. This can be understood through the lens of graph theory and the foundational work of Carothers^44^, Flory^39^, and Stockmayer^45^ on critical gelation point. The degree of a node (individual polymer chain) in a network indicates how many edges (crosslinks) it has^46^. A network is said to be connected when every pair of nodes is connected by a path of arbitrary length, analogous to Carothers’ interpretation of the critical gelation point as the emergence of a polymer with infinite length^44^. As a network’s degree distribution skews right (polymer f increases), achieving connectedness (gelation) occurs with fewer edges (crosslinks) required. Networks with a right-skewed degree distribution have many redundant paths between nodes, enabling robustness to disruption.

The impact of polymer f on network structure is visualized in **Fig. 3a**, where the topology of step-growth polymerized hydrogels with equivalent functional group concentrations but increasing f (decreasing polymer molecule count) is modeled. At f = 4, polymer chains can be connected to at most 4 other chains. The network becomes connected only when the vast majority of possible crosslinks are formed. As f increases, network degree distribution skews right, drastically decreasing the number of crosslinks needed for gelation. The decreasing ratio of critical crosslinks to all crosslinks as f increases in **Fig. 3a** illustrates this. Conversely, the number of crosslinks that need to be disrupted to disconnect the fully formed network increases with f.

**Figure 3.**
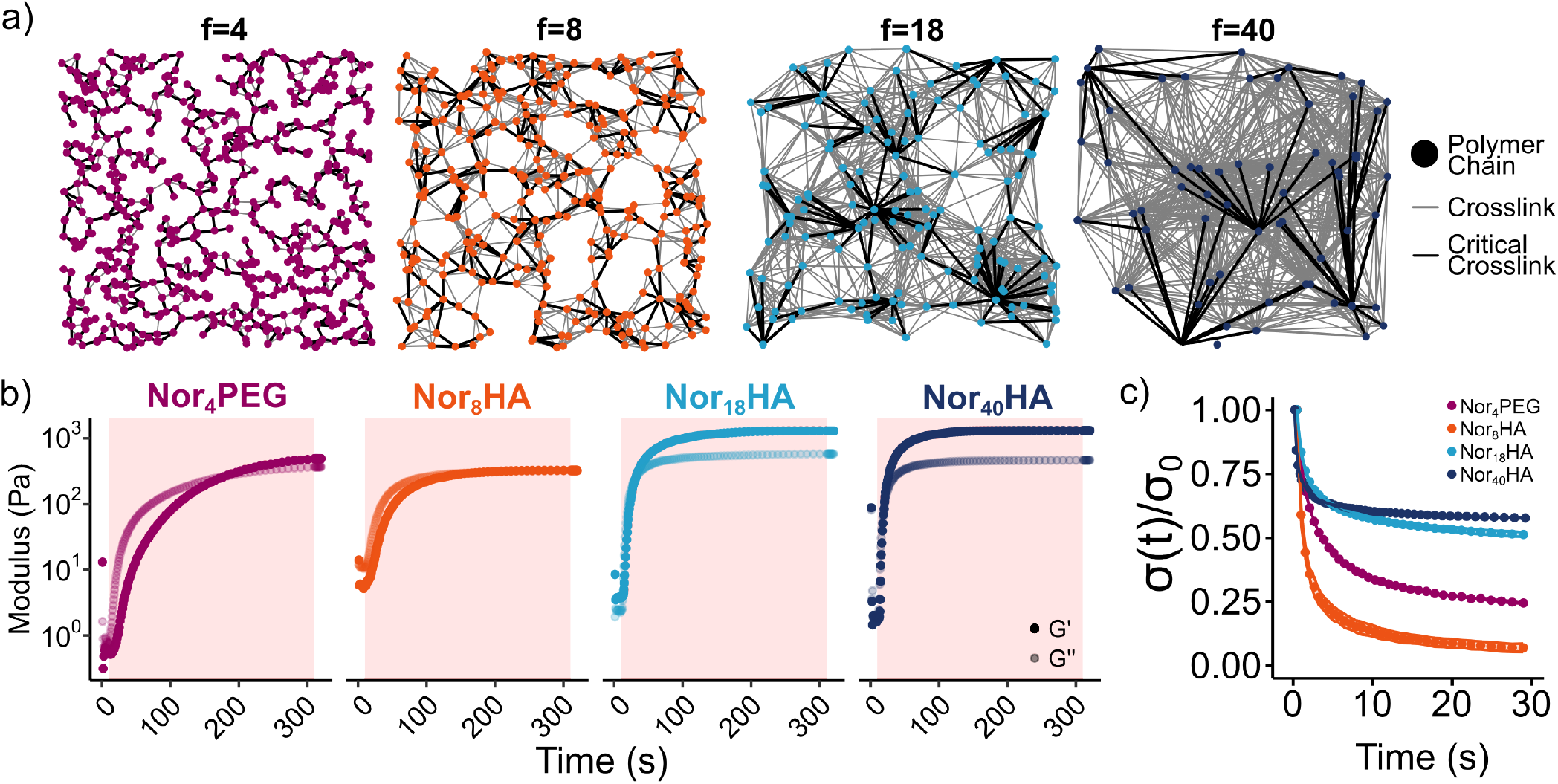
Modifying NorHA f to circumvent norbornene homopolymerization-driven network formation. a) Representations of network structure in fully step-growth crosslinked hydrogels of various f. Each node is an individual polymer chain, and connections between nodes are crosslinks. Critical crosslinks are a minimal set of crosslinks needed to connect all polymer chains. As f increases, the proportion of critical crosslinks drastically decreases, illustrating that network formation readily occurs even at low conversion of highly modified polymers. b) To determine if lowering NorHA f improved stress relaxation extent in the photocurable guest-host system, hydrogel precursor solutions containing one of the norbornene-modified polymers, CDHA, and thiolated adamantane were exposed to UV light for 5 minutes. Polymer and peptide concentration were set so that the molar ratios of norbornene, thiol, adamantane, and β-cyclodextrin were equal. c) Stress relaxation (stress over time, σ(t), normalized to initial stress, σ(0)) tests demonstrated incomplete relaxation of Nor_18_HA and Nor_40_HA hydrogels. Nor_8_HA hydrogels exhibited full stress relaxation, mimicking the behavior of the spontaneously gelling system shown in Fig. 1d.

Carothers first used this graph theoretical approach to describe gelation, and Flory and Stockmayer later provided additional refinement. The functional group conversion at critical gelation point (α^c^) is predicted by Flory-Stockmayer theory to be inversely proportional to f:

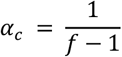

This theory assumes that polymerization is a step-growth process and network loops are nonexistent. We demonstrated that the thiol-norbornene reaction is a hybrid step/chain-growth polymerization. Loops are prevalent in dilute systems as the next closest functional group is often on the same polymer^47^. Flory-Stockmayer theory was demonstrated to not accurately predict gel point of chain-growth networks^48^. On the other hand, the MALDI-ToF spectra showed a maximum chain length of 2 (**Fig. 2c**), indicating that norbornene homopolymerization may be viewed as a pseudo-step growth reaction in this context.

While we cannot accurately predict the critical gelation point of this system using Flory-Stockmayer theory to select an appropriate polymer f, the general concept can still be applied. We hypothesized that decreasing f should facilitate increased hydrogel stress relaxation and encapsulated cell spreading through two avenues: covalently crosslinked network formation can be suppressed even at high rates of norbornene homopolymerization; and decreased network stability will allow for more efficient network remodeling as fewer crosslinks need to be disrupted.

We made a series of polymers with f lower than the previously used NorHA (f = 47) that demonstrated limited stress relaxation capability. We synthesized NorHA of different molecular weights but similar mean degree of modification to maintain norbornene concentration, highlighting that differences in network topology drive these changes, not the norbornene homopolymerization itself. Mean modification was quantified from ^1^H NMR spectra (**Fig. S11**): 19 kDa NorHA was 17% modified for a mean f of 8 (Nor_8_HA), 37 kDa NorHA was 19% modified for a mean f of 18 (Nor_18_HA), and 82 kDa NorHA was 19.5% modified for a mean f of 40 (Nor_40_HA).

To validate that f could influence network formation by norbornene homopolymerization, oscillatory rheological testing was used to monitor gelation kinetics during UV curing of these different polymers (**Fig. S12**). Norbornene concentration and LAP concentration (8.25 mM and 4 mM respectively) were maintained across all groups and increasing amounts of cysteine were included to determine if available thiols inhibited network formation. The Nor_8_HA did not yield a hydrogel even at 0 mM cysteine concentration. The higher f polymers both gelled in the absence of thiol, with Nor_40_HA yielding a stiffer hydrogel than Nor_18_HA as expected (G’ of 592±17.8 Pa and 201±21.9 Pa respectively). Storage modulus decreased at 0.25x cysteine, and no hydrogels were formed at 0.5x cysteine concentration regardless of f.

With f-dependent network formation verified, we fabricated photocurable guest-host hydrogels with the polymers of different f to determine if any of the hydrogels could fully stress relax (**Fig. 3b**). Hydrogel precursor solutions containing one of the norbornene-modified polymers, CDHA, thiolated adamantane, and LAP were exposed to UV light for 5 minutes. Polymer and peptide concentration were set so that the molar ratios of norbornene, thiol, adamantane, and β-cyclodextrin were equal. The Nor_8_HA group behaved similarly to the spontaneously gelling system (Fig. 1a) with a loss factor close to 1 and similar G’ and G’’. Stress relaxation tests showed the desired full relaxation in the Nor_8_HA group, with less relaxation achieved by the higher f polymers (**Fig. 3c**). The Nor_18_HA and Nor_40_HA groups both exhibited limited stress relaxation as seen with the previously used higher f NorHA (Fig. 1d).

We next wanted to determine the extent to which norbornene homopolymerization impacts cell spreading in 3D environments. Hydrogels crosslinked solely by guest-host interactions do not possess long-term stability, so a dithiol MMP-degradable peptide crosslinker was added to the system along with thiolated RGD adhesive ligand to support cell attachment. Modes of cell-driven remodeling of this substrate are physical deformation via disruption of guest-host interactions and enzymatic scission of the MMP-degradable peptide. Non-degradable covalent crosslinks introduced by norbornene homopolymerization could inhibit remodeling (and therefore, cell spreading) via these processes.

Human mesenchymal stromal cells (hMSCs) were cultured in hydrogels with identical formulations but with different polymers of varying f. A Nor_4_PEG group was included as a positive control for expected cell behavior in networks that are easy to remodel. Concentrations of all reagents including crosslinkers were maintained across all groups, theoretically maintaining norbornene homopolymerization rate across all groups. Only network topology should differ between groups. Note that these hydrogels contained a thiol-to-norbornene molar ratio of 0.87:1, which is well within the range that should not over inflate norbornene homopolymerization due to limited thiol availability. Hydrogel mechanical testing suggested differences in network topology, as G’ increased from 0.7 kPa to 4.5 kPa as f increased (*p* = 7.24×10^-7^, **Figs. 4a and 4b**). Similarly, the Nor_4_PEG and Nor_8_HA groups exhibited a much higher extent of stress relaxation compared to that of the Nor_18_HA and Nor_40_HA groups (*p* = 2.98×10^-11^), as seen in **Fig. 4c**.

**Figure 4.**
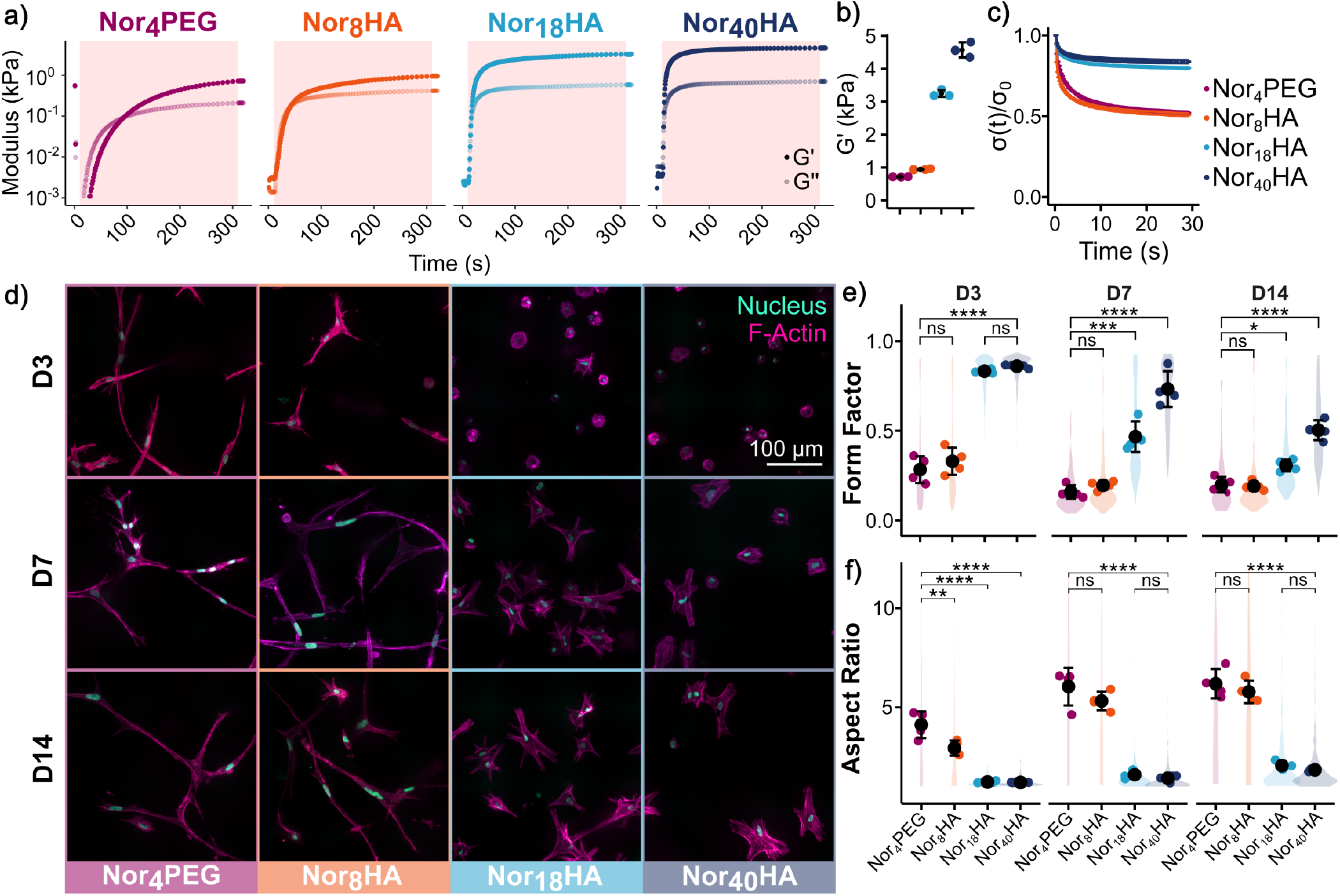
Higher polymer f impedes cell spreading. a) To determine if polymer f impacted encapsulated cell spreading, hydrogel precursor solutions containing one of the norbornene-modified polymers, CDHA, thiolated adamantane, MMP-degradable crosslinker, and LAP were exposed to UV light for 5 minutes. The only difference in formulation between groups was norbornene-modified polymer concentration to ensure constant norbornene concentration. b) As expected, final storage modulus (G’) increased as polymer f increased. c) The low-f polymer hydrogels exhibited stress relaxation extent close to 50%, while the high-f polymer hydrogels only relaxed to about 20%. No group exhibited complete stress relaxation due to inclusion of covalent MMP-degradable crosslinks. d) Z-projected confocal microscopy images of encapsulated hMSCs after 3, 7, and 14 days of culture. Nuclei are shown in cyan (DAPI), and F-actin is shown in magenta (phalloidin). e) Cell circularity was quantified as form factor, where 1 is a perfect circle, and 0 is highly irregular. The hMSCs within the Nor_4_PEG and Nor_8_HA hydrogels were irregularly shaped throughout the culture period, while those within the Nor_18_HA and Nor_40_HA hydrogels became less circular at each successive time point. f) While hMSCs within all groups were irregularly shaped by day 14, morphological differences between low-f and high-f polymer hydrogels were visually apparent. Quantification of cell aspect ratio demonstrated more elongated morphology in low-f polymer hydrogels. The mean and standard deviation are shown as a single black dot with error bars. The colored dots are the mean metric of each hydrogel. The violin plots show the distribution of values calculated for each cell. Statistical analyses were performed on the hydrogel means via ANOVA followed by post-hoc Tukey’s HSD tests. *N* = 4 hydrogels, 131-704 cells per group. ****: *p* < 1×10^-5^, ***: *p* < 1×10^-3^, **: *p* < 0.01, *: *p* < 0.05.

We monitored cell morphology over two weeks via fluorescent imaging of phalloidin-stained F-actin cytoskeleton (**Fig. 4d**). Extent of cell spreading was quantified through two metrics: form factor, a measure of circularity calculated as 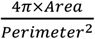, and aspect ratio, calculated by dividing the major axis length by the minor axis length of circumscribing ellipses of each cell. Cell circularity was low in the Nor_4_PEG and Nor_8_HA groups across all time points, and the Nor_18_HA and Nor_40_HA groups became more irregular at each successive time point (**Fig. 4e**). Despite high cell shape irregularity in all groups, morphology was visually very different across groups at day 14. Aspect ratio was used to quantify this difference as the hMSCs within the Nor_4_PEG and Nor_8_HA hydrogels were elongated and those within the Nor_18_HA and Nor_40_HA hydrogels had more compact morphology (**Fig. 4f**). We then used the CAJAL_49_ python package to explore morphological differences not adequately captured by predetermined metrics at a single-cell resolution. CAJAL calculates the Gromov-Wasserstein distance^50^, a translation- and rotation-invariant metric often used in shape-matching applications, between all pairs of cells. This metric measures the cost of transforming one shape, a cell mask in this context, into another, with dissimilarly shaped objects being further away from one another. We then clustered the resultant distance matrix with the Leiden clustering algorithm, yielding 29 subpopulations with obvious differences in size and shape. The distance matrix is then embedded into a lower dimensionality for visualization using the UMAP algorithm to yield a 2D morphological space (**Fig. 5a**). Representative cell masks of each subcluster are placed adjacent to their respective subcluster to visualize the diverse morphologies identified.

**Figure 5.**
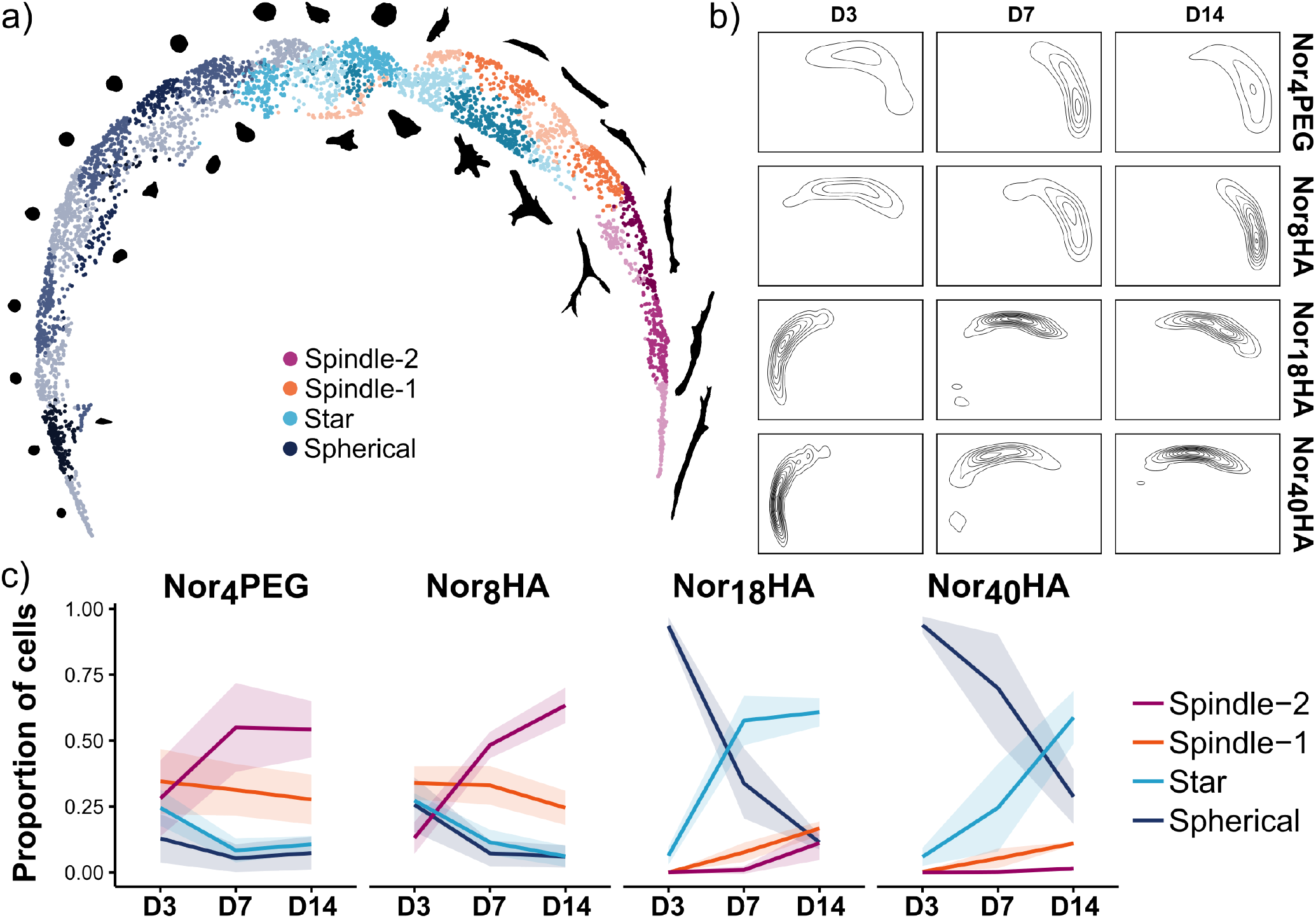
A latent morphological space reveals two distinct cell shape trajectories associated with either low-f polymer hydrogels or high-f polymer hydrogels. a) A 2D embedding of the morphological space inferred by CAJAL. Each point is one of 4509 cells, and points are colored according to broad morphological category. Different shades of the four primary colors indicate different subclusters that were grouped together to form the morphological categories. Masks of representative cells belonging to these subclusters are positioned adjacent to their corresponding subcluster, illustrating the changes in morphology identified by CAJAL. b) Contour maps showing density of points throughout the morphological space. Maps are split according to day (columns) and polymer (rows) to visualize the differences in cell shape transitions between polymers. c) Cell shape dynamics across day 3, day 7, and day 14 were quantified by tabulating the proportions of cells within the four morphological categories for each polymer. The midline represents the mean proportion calculated from *n* = 4 hydrogels, and the ribbon indicates the standard deviation. The Nor_4_PEG and Nor_8_HA groups show slight decreases in Spherical, Star, and Spindle-1 cells with a concomitant increase in more elongated Spindle-2 cells. Within the Nor_8_HA and Nor_18_HA groups, the primary dynamic was the transition from Spherical to Star morphology.

Two broad groups were apparent: a group of spindle morphology cells primarily composed of cells from Nor_4_PEG and Nor_8_HA hydrogels, and another group showing a trajectory from small spherical morphology to a larger starshaped morphology primarily composed of cells from Nor_18_HA and Nor_40_HA hydrogels. To simplify interpretation of differences in morphological dynamics between conditions, the clusters were manually grouped into four categories based on morphology of the representative cells: Spherical, Star, Spindle-1, and Spindle-2. Population dynamics are visualized as contour maps of each hydrogel condition at each time point (**Fig. 5b**). Cells within the Nor_4_PEG and Nor_8_HA hydrogels start primarily in the Star/Spindle-1 region and transition to primarily Spindle-2 by day 14. Cells within the Nor_18_HA and Nor_40_HA hydrogels start primarily in the Spherical region and transition to primarily Star by day 14. To more robustly quantify this transition, proportions of cells within each morphological category were tabulated at each time point for each hydrogel condition (**Fig. 5c**). The hMSCs within Nor_4_PEG and Nor_8_HA hydrogels transitioned toward an elongated Spindle-2 morphology with time, as Spherical and Star morphology cells decreased. The Nor_18_HA and Nor_40_HA hydrogels restricted spreading as nearly all hMSCs were Spherical at day 3. Over time, cells primarily transitioned to the Star morphology with few Spindle cells.

The stark morphological differences between cells within low-f and high-f hydrogels can be jointly attributed to differences in substrate remodeling capability and stiffness. Stiffness is a well-established regulator of cell behavior in 3D^22,51^; very soft hydrogels fail to activate the cytoskeletal machinery necessary for adhesion and remodeling of networks, yielding a round quiescent phenotype^52^. High stiffness hydrogels result in compact star-like cell morphology due to inefficient remodeling of dense networks. Intermediate stiffness hydrogels facilitate optimal spreading as mechanotransduction pathways are activated and matrix remodeling is not inhibited by dense crosslinking. Matrix degradation has also been shown to modify kinetics of spreading, but not final morphology^51^. If these cultures were prolonged, it is possible cells within the high-f polymer hydrogels would have similar spindle morphology as the low-f polymer hydrogels.

### Norbornene homopolymerization impedes cell spreading in stiffness-matched hydrogels

The purpose of the previous experiment was to demonstrate how network topology differences caused by polymer f could inhibit cell spreading, specifically from norbornene homopolymerization forming non-degradable static crosslinks. However, hydrogel stiffness varied greatly across groups, muddying the interpretation as substrate stiffness and viscoelasticity both influence cell behavior. Therefore, we next sought to control for this by encapsulating hMSCs in hydrogels of variable f but equivalent mechanics.

A Nor_40_HA guest-host supramolecular hydrogel without MMP-degradable crosslinks was first fabricated, and the hydrogels made from the other polymers were fabricated to have similar G’ and G’’ by changing guest-host concentration, MMP-degradable crosslinker concentration, and polymer concentration (**Fig. 6a**). *In situ* mechanics were similar across all groups (**Fig. S13**). These hydrogels were allowed to swell overnight in PBS, and further mechanical testing demonstrated similar G’ in all but the Nor_4_PEG group (*p* = 7.63×10^-6^, **Fig. 6b**), and similar stress relaxation behavior across all groups (*p* = 0.280, **Fig. 6c**). To demonstrate differences in their ability to be remodeled, preswollen hydrogels were incubated in 150 U mL^-1^ collagenase. After just 12 hours, the Nor_4_PEG and Nor_8_HA hydrogels were completely degraded (**Fig. S14**). As expected, the Nor_40_HA hydrogels did not appear to degrade at all. Despite having MMP-degradable crosslinks, the Nor_18_HA hydrogels remained stable, even in this high concentration of collagenase.

**Fig. 6.**
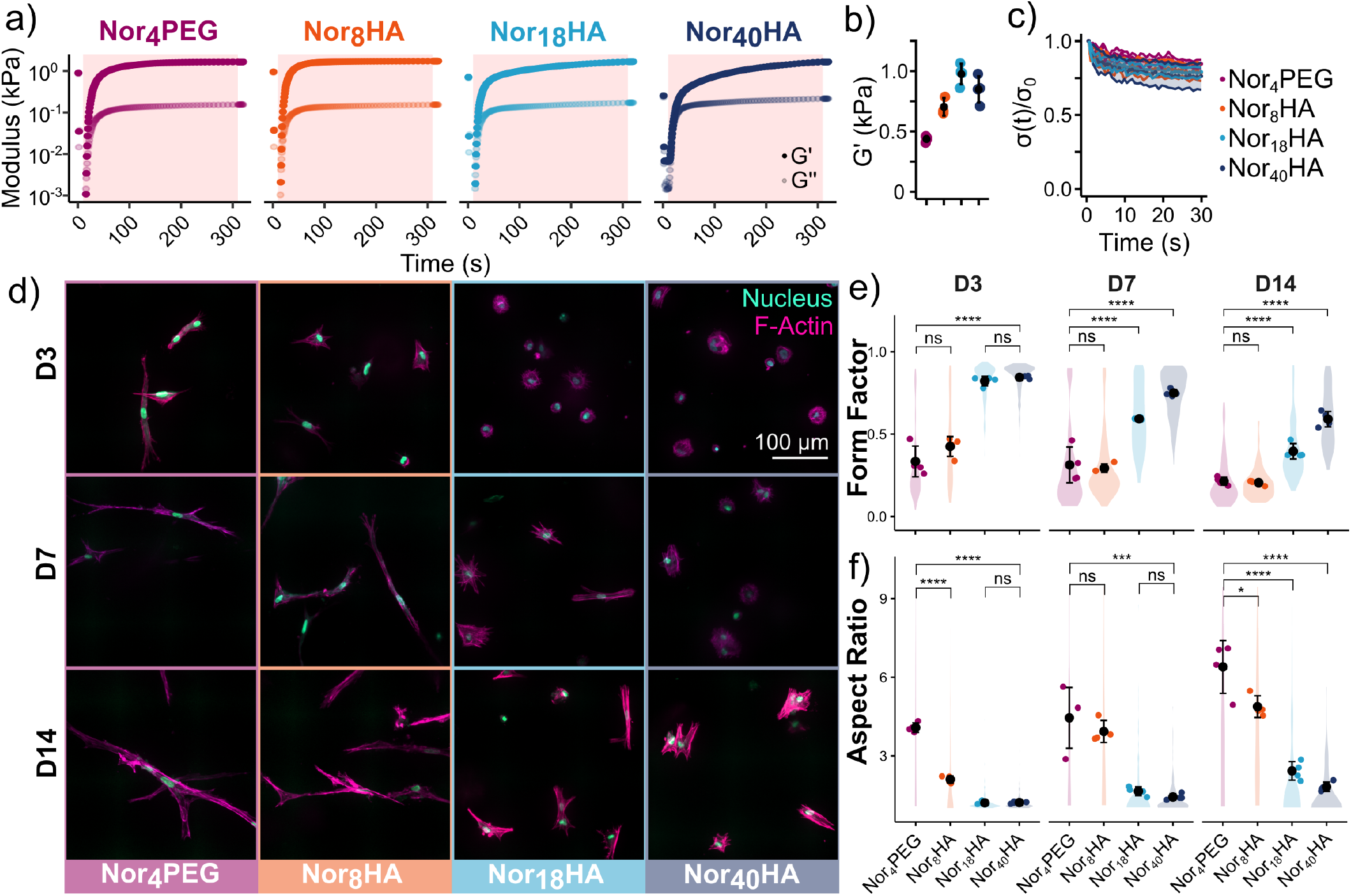
Norbornene homopolymerization, not substrate mechanics, alters cell morphology. a) To determine if matrix mechanics drove the differences in cell morphology attributed to polymer f in Figs. 4 and 5, we created a series of hydrogels with variable f but equivalent mechanics. Hydrogel precursor solutions containing one of the norbornene-modified polymers, CDHA, thiolated adamantane, LAP, and MMP-degradable crosslinker at various concentrations were exposed to UV light for 5 minutes. The Nor_40_HA hydrogel did not contain any MMP-degradable crosslinker; crosslinker was added to the other formulations to reach the stiffness of the Nor_40_HA hydrogels. b) Final storage modulus (G’) was around 900 Pa for the NorHA polymer hydrogels. The PEG polymer hydrogel was considerably softer but within the same order of magnitude. c) All hydrogels exhibited stress relaxation extent close to 20%. d) Z-projected confocal microscopy images of encapsulated hMSCs after 3, 7, and 14 days of culture. Nuclei are shown in cyan (DAPI), and F-actin is shown in magenta (phalloidin). e) The hMSCs within the Nor_4_PEG and Nor_8_HA hydrogels were irregularly shaped throughout the culture period, while those within the Nor_18_HA and Nor_40_HA hydrogels became less circular at each successive time point. f) Quantification of cell aspect ratio demonstrated more elongated morphology in low-f polymer hydrogels compared to high-f polymer hydrogels at all time points. The mean and standard deviation are shown as a single black dot with error bars. The colored dots are the mean metric of each hydrogel. The violin plots show the distribution of values calculated for each cell. Statistical analyses were performed on the hydrogel means via ANOVA followed by post-hoc Tukey’s HSD tests. *N* = 4 hydrogels, 79-256 cells per group. ****: *p* < 1×10^-5^, ***: *p* < 1×10^-3^, **: *p* < 0.01, *: *p* < 0.05.

The hMSCs encapsulated in these hydrogels showed similar changes in morphology as seen in the previous experiment. Cell circularity was low in the Nor_4_PEG and Nor_8_HA groups across all time points, and the Nor_18_HA and Nor_40_HA groups became more irregular at each successive time point (**Fig. 6e**). By two weeks, form factor was similarly low across all hydrogels but the Nor_40_HA group. Aspect ratio again showed that hMSCs in the Nor_4_PEG and Nor_8_HA groups had significantly more elongated morphology than those in the Nor_18_HA and Nor_40_HA groups at all time points (**Fig. 6f**).

The morphological space inferred by CAJAL for this experiment was similar to the previous encapsulation. The hMSCs within the Nor_4_PEG and Nor_8_HA hydrogels primarily fell within a group of cells with spindle morphology, while cells encapsulated in the Nor_18_HA and Nor_40_HA hydrogels fell along a spherical-to-star trajectory (**Fig. 7a**). Population dynamics were visualized as contour maps of each hydrogel condition at each time point (**Fig. 7b**). Cells within the Nor_4_PEG and Nor_8_HA hydrogels were primarily elongated and the presence of spherical cells diminished over time. Cells within the Nor_18_HA and Nor_40_HA hydrogels transitioned from primarily spherical morphology to primarily star morphology by day 14.

**Figure 7.**
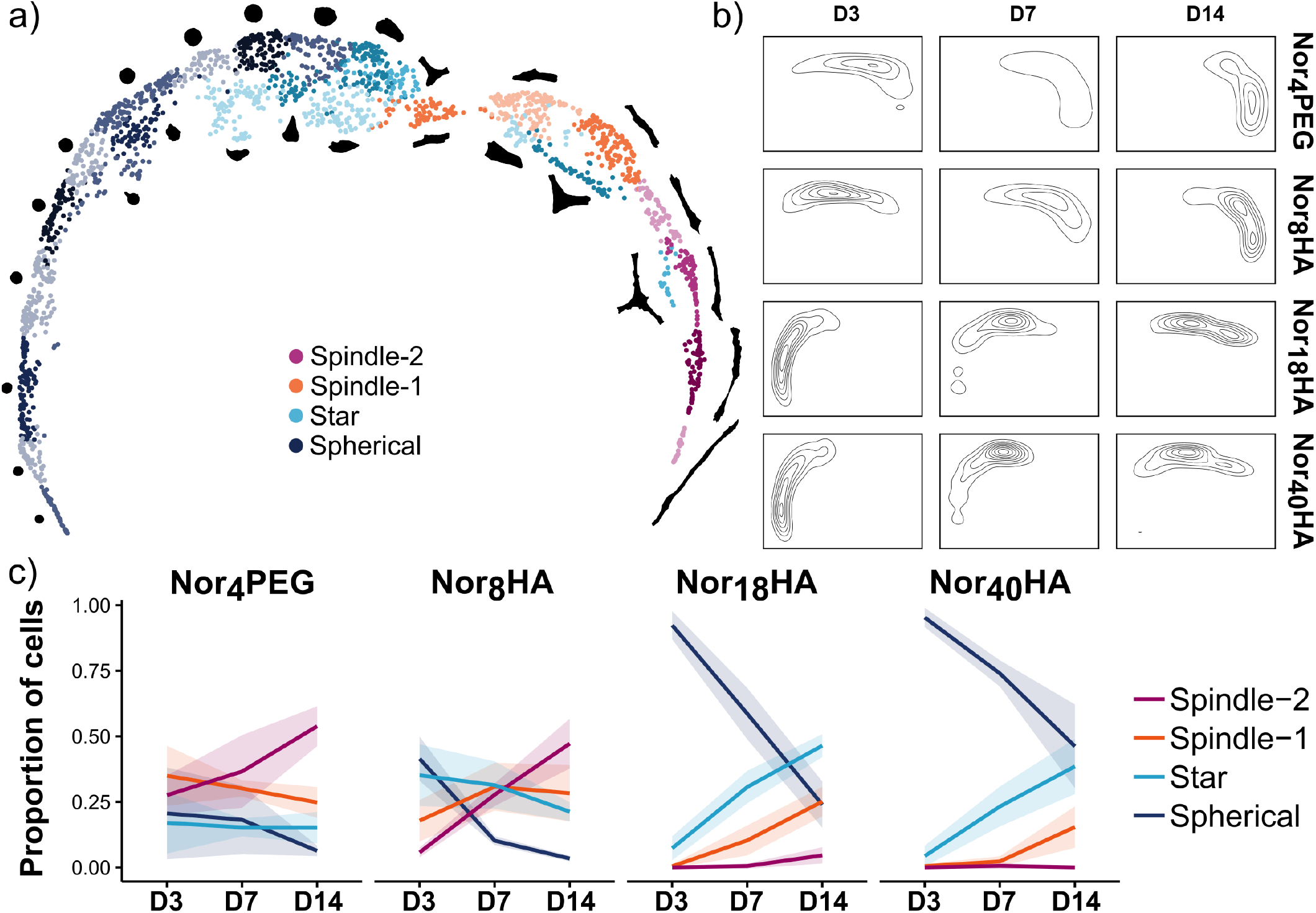
The two distinct cell shape trajectories reappeared in mechanically-matched hydrogels. a) A 2D embedding of the morphological space inferred by CAJAL. Each point is one of 2069 cells, and points are colored according to broad morphological category. Different shades of the four primary colors indicate different subclusters that were grouped together to form the morphological categories. Masks of representative cells belonging to these subclusters are positioned adjacent to their corresponding subcluster, illustrating the changes in morphology identified by CAJAL. b) Contour maps showing density of points throughout the morphological space. Maps are split according to day (columns) and polymer (rows) to visualize the differences in cell shape transitions between polymers. c) Cell shape dynamics across day 3, day 7, and day 14 were quantified by tabulating the proportions of cells within the four morphological categories for each polymer. The midline represents the mean proportion calculated from *n* = 4 hydrogels, and the ribbon indicates the standard deviation. The Nor_4_PEG and Nor_8_HA groups showed slight decreases in Spherical, Star, and Spindle-1 cells with a concomitant increase in more elongated Spindle-2 cells. Within the Nor_8_HA and Nor_18_HA groups, the primary dynamic was the transition from Spherical to Star morphology.

Quantification of morphological category kinetics showed that hMSCs within Nor_4_PEG and Nor_8_HA hydrogels transitioned toward an elongated Spindle-2 morphology with time, as Spherical and Star morphology cells decreased (**Fig. 7c**). The Nor_18_HA and Nor_40_HA hydrogels restricted spreading as nearly all hMSCs were Spherical at day 3. Over time, cells primarily transitioned to the Star morphology with few Spindle cells.

## CONCLUSIONS

Unexpected partial stress relaxation of a radically initiated physically-crosslinked thiol-norbornene hydrogel led us to investigate the prevalence of norbornene homopolymerization in these supposed click reactions. Norbornene conversion exceeded input thiol concentration across all tested system permutations, verifying the presence of norbornene homopolymerization regardless of thiol concentration. These findings call into question the classification of radical-initiated thiol-norbornene polymerization as a click reaction in dilute aqueous conditions used to make hydrogels. Norbornene homopolymerization affects any system that uses radical-initiated thiol-norbornene polymerization, though hydrogel performance may be less impacted when using lower f polymers.

Instead of attempting to mitigate norbornene homopolymerization, we circumvented its effects by reducing the degree of functionalization on our polymer from approximately 45 to 8. Lowering polymer f rescued the desired stress relaxation behavior in our guest-host hydrogel system. The impact of norbornene homopolymerization and polymer f on cell spreading was assessed by encapsulating hMSCs in hydrogels of same formulation but varying f. Cells within the low-f polymer hydrogels appeared to easily remodel the matrix as they had an elongated, spindle-like morphology. The high-f polymer hydrogels restricted cell spreading, as cells were primarily rounded or exhibited a compact star morphology. To account for the different stiffnesses of these matrices, mechanics were then held constant by primarily changing concentration of MMP-degradable crosslinker. The same spindle vs. star morphological divergence between low-f and high-f polymer hydrogels occurred, suggesting that mechanics did not drive this phenotypic difference. Norbornene homopolymerization was shown to introduce undesirable non-degradable covalent crosslinks that impede encapsulated cell spreading. This limitation was overcome by fabricating networks from lower f polymers with reduced connectivity, demonstrating the importance of considering network properties in addition to crosslink mechanism when designing engineered cellular microenvironments.

## EXPERIMENTAL METHODS

### Materials

Sodium hyaluronate (sodium HA, 19, 37, or 82 kDa) was purchased from Lifecore Biosciences. β-Cyclodextrin (β-CD), hexamethylenediamine (HDA), ammonium chloride, and p-toluenesulfonyl chloride were purchased from TCI America. Tetrabutylammonium hydroxide (TBA-OH) was purchased from Acros Organics. NorPEG (4-arm, 20 kDa) was purchased from Biopharma PEG. Methyl-PEG-norbornene (1 kDa) was purchased from Creative PEGWorks. All other materials were purchased from Milli-poreSigma-Aldrich unless stated otherwise.

### Synthesis of hyaluronic acid tert-butyl ammonium salt (HA-TBA)

Hyaluronic acid tert-butyl ammonium salt was synthesized as previously reported^40^. Briefly, sodium hyaluronate (Lifecore, 82 kDa) was reacted with Dowex 50W proton-exchange resin, filtered, titrated with TBA-OH to pH 7.05, frozen, and lyophilized to yield HA-TBA.

### Synthesis of cyclodextrin-modified hyaluronate (CDHA)

An amine functionalized form of β-CD, 6-(6-aminohexyl)amino-6-deoxy-β-cyclodextrin (β-CD-HDA), was first synthesized for subsequent amide coupling with HA-TBA. As described previously, p-toluenesulfonyl chloride was added dropwise to aqueous β-cyclodextrin (β-CD) at 0°C to form 6-O-monotosyl-6-deoxy-β-cyclodextrin (CD-Tos)^40^. After product confirmation with ^1^H NMR, CD-Tos was reacted with hexamethylenediamine in DMF for 18 h at 80°C under nitrogen to form β-CD-HDA.

CDHA was prepared via (benzotriazol-1-yloxy) tris(dimethylamino) phosphonium hexafluorophosphate (BOP)-mediated amide coupling between HA-TBA and β-CD-HDA. HA-TBA, β-CD-HDA, and BOP were reacted in DMSO for 3 h at 25°C. The reaction was quenched with cold water, dialyzed against water, sterile filtered, frozen, and lyophilized. The degree of modification was 22% as determined by ^1^H NMR (**Fig. S1**).

### Synthesis of adamantane-modified hyaluronate (AdHA)

AdHA was prepared via di-tert-butyl bicarbonate (BOC_2_O)/4-dimethylaminopyridine (DMAP)-mediated esterification between HA-TBA and 1-adamantaneacetic acid^40^. HA-TBA, adamantaneacetic acid, BOC^2^O, and DMAP were reacted in DMSO for 24 h at 45°C. The reaction was quenched with cold water, dialyzed against water, sterile filtered, frozen, and lyophilized. The degree of modification was 26% as determined by ^1^H NMR (**Fig. S2**).

### Synthesis of norbornene-modified hyaluronate (NorHA)

NorHA was prepared via 4-(4,6-dimethoxy-1,3,5-triazin-2-yl)-4-methylmorpholinium chloride (DMTMM)-mediated amide coupling between sodium hyaluronate (Lifecore, 82 kDa) and 5-norbornene-2-methylamine^53^. HA was dissolved in aqueous MES buffer at pH of 5.5. Excess DMTMM was added to the solution and stirred for 15 min. 5-norbornene-2-methylamine was added dropwise to the solution, which was subsequently allowed to react at room temp overnight. The solution’s pH was adjusted to 11 using NaOH for 15 min to dissolve precipitates formed by high DMTMM concentrations. The reaction was precipitated in cold ethanol, dialyzed against water, sterile filtered, and lyophilized. The degree of modification was 24% as determined by ^1^H NMR (**Fig. S3**).

The same procedure was used for the three additional NorHA polymers synthesized for the f comparison studies (19 kDa, 37 kDa, and 82 kDa respectively). The degrees of modification were 17%, 18.5%, and 19% respectively as determined by ^1^H NMR (**Fig. S12**).

### Synthesis of thiolated adamantane peptide

Solidphase peptide synthesis of the Ad-KKKCG peptide was performed using an automated, microwave-assisted Liberty Blue peptide synthesizer (CEM) on Rink Amide ProTide^TM^ resin (CEM, 0.65 mmol g^-1^)^33^. Fmoc-protected amino acids (Fmoc-Lys(Boc)–OH, Fmoc-Gly–OH, Fmoc-Cys(Trt)–OH from Gyros Protein Technologies) and 1-adamantaneacetic acid were dissolved in DMF at a concentration of 0.2 M and added to the reaction vessel according to the peptide sequence, with deprotection and coupling steps between additions. Piperidine in DMF (20% v/v) was the Fmoc deprotection agent, and coupling was carried out at 90°C for 2 min using diisopropylcarbodiimide (1 M in DMF) and Oxyma Pure (1 M in DMF).

Peptide cleavage and deprotection of chain side groups was carried out in a cleavage cocktail containing 92.5% (v/v) trifluoroacetic acid, 2.5% triisopropylsilane, 2.5% 2,2′-(ethylenedioxy)diethanethiol, and 2.5% ultrapure water for 2 h at room temperature. The peptide was precipitated in cold ether, isolated via centrifugation, and underwent another cold ether wash and isolation step. The precipitate was dried under vacuum, dissolved in water, frozen, and lyophilized. Synthesis of the desired product of high purity was confirmed via MALDI-ToF after reduction via TCEP (**Fig. S4**).

### Proton nuclear magnetic resonance

^1^H NMR experiments were performed on a Bruker Neo 400 MHz NMR Spectrometer. Pure polymer samples were dissolved in D_2_O at 20 mg mL^-1^. For the conversion studies, NorHA samples that gelled were first degraded with hyaluronidase. All samples were then lyophilized and resuspended in D_2_O.

### Matrix-assisted laser desorption/ionization mass spectrometry

MALDI mass spectrometry experiments were performed in linear mode at 25% laser power with a 355 nm, 200 Hz solid-state laser (MALDI-8020, Shimadzu) using a α-cyano-4-hydroxycinnamic acid matrix (5 mg mL^-1^ in acetonitrile). Samples were diluted in acetonitrile to approximately 1 µM. Matrix solution was deposited in 0.5 µL drops on the sample plate wells and mixed with 0.5 µL sample solution prior to air drying.

### Fabrication of spontaneously crosslinked guest-host hydrogels

Concentrated stock solutions of CDHA and AdHA were mixed with PBS to yield guest-host hydrogels with 1:1 ratio of cyclodextrin:adamantane groups (32.5 mg mL^-1^ of 22% modified CDHA yielding 11 mM β-cyclodextrin, 18 mg mL^-1^ of 26% modified AdHA yielding 11 mM adamantane).

### Fabrication of radically crosslinked guest-host hydrogels

Thiolated adamantane peptide (Ad-KKK**C**G) and MMP-degradable crosslinker (GenScript, G**C**NS-GPQG↓IWGQ-SN**C**G) were treated with TCEP (pH 7.0) at a 0.5:1 molar equivalent of TCEP:thiol prior to preparation of hydrogel precursor. Concentrated stock solutions of CDHA and thiolated adamantane were first mixed together to allow complexation. NorHA, LAP (4 mM final concentration), and PBS (enough to reach final desired volume) were added to the CDHA/thiolated adamantane mixture prior to vortexing. Photocurable hydrogels were fabricated by exposing precursor solutions to 5 mW cm^-2^ 365 nm light using an Omnicure Series 1500 UV lamp (Excelitas).

For the spontaneous versus photocurable comparison in Fig. 1, a hydrogel precursor solution mimicking the formulation of the spontaneous hydrogel was prepared by maintaining the same concentration of CDHA and using enough NorHA for a 1:1 molar ratio of norbornene:cyclodextrin. Adamantane peptide was added at a 1:1 molar ratio to norbornene (32.5 mg mL^-1^ of 22% modified CDHA yielding 11 mM β-cyclodextrin, 20 mg mL^-1^ of 24% modified NorHA yielding 11 mM norbornene). For the NorPEG samples, polymer was at a concentration such that norbornene concentration matched the NorHA-based samples.

### Rheological characterization

All rheological measurements were performed at 25°C on an Anton Paar MCR 302 using either a cone plate (25 mm diameter, 0.5°, 25 µm gap) for most experiments, or with a parallel plate (8 mm diameter) for measurements on swollen hydrogels. To monitor mechanics during UV curing, oscillatory time sweeps (1% strain, 1 Hz) were used with UV irradiation (365 nm, 5 mW cm^-2^) for 5 min. For stress relaxation tests, shear stress was monitored over 30 sec after application of 10% strain.

### Degradation studies

Hydrogels (*n* = 4, 30 µL each) were incubated overnight in 1 mL solution of 150 U mL^-1^ Collagenase, 1x PBS without CaCl_2_, 5 mM CaCl_2_, and 0.1% Triton X-100.

### Norbornene conversion quantification

Hydrogel precursor solutions contained enough polymer to yield 11.0 mM norbornene (20 mg mL^-1^ for 22% modified NorHA and 41.25 mg mL^-1^ for NorPEG); thiol at 0 mM (0.0x), 2.2 mM (0.2x), 4.4 mM (0.4x), 6.6 mM (0.6x), 8.8 mM (0.8x), or 11.0 mM (1.0x); and 4 mM LAP in PBS. Thiols were reduced with 0.5:1 molar ratio of TCEP:thiol prior to mixing. Hydrogel precursors were exposed to 365 nm light at 5 mW cm^-2^ for 5 min. Samples made with NorHA were then digested in 0.3 mg mL^-1^ hyaluronidase and 0.1% Triton X-100 at 37°C overnight. All samples were flash frozen and subsequently lyophilized before being dissolved in deuterated water for ^1^H NMR.

### Network representations of crosslinked networks of varying polymer f

Network representations were created with the python package NetworkX and visualized using the python package matplotlib. Nodes were placed randomly in a 2D space and connected to their n nearest neighbors. Nodes could only crosslinks at most once to another node. For the f = 4 network, nodes were connected to exactly 4 other nodes (n = 4) to mimic crosslinking of Nor_4_PEG. For the other networks, nodes were connected on average n = f nodes, with n following a Poisson distribution as described in the previous paragraph. This mimicked crosslinking of a randomly-modified polymer like NorHA. The minimum spanning tree algorithm was used to identify the minimal set of crosslinks needed to link all polymer chains, which corresponds to Carothers’ interpretation of minimal network formation^44^. Critical crosslink count closely followed Carothers’ definition of critical gelation point (p_c_ = 2/f). Code used to create these networks can be viewed on GitHub (see Code Availability).

### Cell encapsulation in hydrogels

Human mesenchymal stromal cells (hMSCs, Lonza) were used at passage 3 for all experiments. Culture media contained Gibco minimum essential medium (MEM-α) supplemented with 20% v/v fetal bovine serum (Gibco) and 1% v/v penicillin/streptomycin/amphotericin B (100 U mL^-1^, 100 μg mL^-1^, and 0.25 μg mL^-1^ final concentrations, respectively, Gibco). Prior to encapsulation, lyophilized materials were dissolved in sterile PBS. Hydrogel precursor solutions were mixed as described earlier, with the addition of thiolated RGD peptide (G**C**GYGRGDSPG, Genscript) at a concentration of 1 mM and hMSCs at 1-2×10^5^ cells per mL. Hydrogel precursor solution was pipetted into round silicone molds (6 mm diameter, 1 mm height) and exposed to UV for 5 min to form 30 µL hydrogel plugs. Media was replaced every 2–3 days.

### Fluorescent staining and confocal imaging

Samples were fixed with 10% formalin in PBS for 15 min. Blocking and permeabilization were performed using 1% BSA and 0.1% Triton X-100 overnight at 4°C. Samples were then incubated overnight at 4°C in DAPI (1:10,000 dilution) and rhodamine-phalloidin (1:2,000 dilution). Imaging was conducted using a Cytation C10 spinning disc confocal microscope (Agilent BioTek) equipped with a 20× objective and a 60 µm pinhole aperture disc. The focal plane was set 50 µm into the hydrogel before capturing 25 z-stacks at 8.4 µm intervals, yielding a total imaging depth of approximately 200 µm.

### Cell segmentation with CellPose

3D image sets were z-projected to yield a 2D representation of each field of view. Using the CellPose 2.0^54^ model ‘cyto2’ implemented in the Omnipose^55^ v1.0.7 python package, cells were segmented using both nuclear (DAPI) and F-actin (rhodamine phalloidin) channels. Specific model parameters can be seen at GitHub (see Code Availability). Masks were then manually corrected for later use with CAJAL. Cell morphological metrics were calculated using the regionprops() function of the scikit-image v0.24.0 python package.

### Cell morphological space using CAJAL

Corrected masks were imported into CAJAL^49^ v1.0.3 with minor file I/O modifications for easier use. Masks smaller than 50 µm^2^ were discarded. First, the intracellular distance matrices for all cells in each experiment were calculated. Then, the Gromov-Wasserstein distance was calculated between all pairs of cells. Similar cell populations were identified via Leiden clustering of the Gromov-Wasserstein distance matrix. Exemplar cells from each cluster were identified using the utilities.identify_medoid() function. A 2-dimensional UMAP representation of the distance matrix was created with the python package umap-learn. Embeddings were visualized with the R package ggplot2. To further investigate morphological dynamics, the portion of cells within each morphological category was tabulated for each sample and plotted using the R packages dplyr and ggplot2. Code used for this analysis can be seen at GitHub (see Code Availability)

## ABBREVIATIONS

CDHA: β-cyclodextrin-modified hyaluronic acid
AdHA: adamantane-modified hyaluronic acid
NorHA: norbornene-modified hyaluronic acid
NorPEG: 4-arm norbornene-modified polyethylene glycol
LAP: lithium acylphosphinate
G’: shear storage modulus
G’’: shear loss modulus
σ: shear stress
^1^H NMR: proton nuclear magnetic resonance
MALDI-ToF: matrix-assisted laser desorption/ionization time-of-flight
f: number of functional groups on a polymer
MMP: matrix metalloproteinase
hMSCs: human mesenchymal stromal cells.

## ASSOCIATED CONTENT

### Supporting Information

The Supporting Information is available free of charge on the ACS Publications website.

Additional figures of supporting data (PDF)

Test statistics and p-values of ANOVA and pair-wise comparisons for data in Figures 4-7 (TSV)

### Code Availability

Code and corresponding data used to produce figures within this manuscript are available at https://github.com/Caliari-Lab/2025_Gentry_Norbornene_Homopolymerization.

### Image Availability

Confocal ICC images and cell masks will be deposited in the BioImage Archive.

## Author Contributions

J.L.G. and S.R.C. conceived the idea and designed experiments.

J.L.G. conducted experiments and associated analysis. J.L.G. and

S.R.C. wrote the manuscript. S.R.C. acquired funding and supervised the work.

## Notes

The authors declare no competing financial interest.

## ACKNOWLEDGMENT

We thank Prof. Rachel Letteri for helpful discussions and use of her peptide synthesizer. This work was supported by the NSF (CAREER 2036592) and NIH (R01AR078886). The content is solely the responsibility of the authors and does not necessarily represent the official views of the National Institutes of Health.

