## Supplemental Information for "Norbornene homopolymerization limits cell spreading in thiol-ene photoclick hydrogels"

### Supplementary Figures

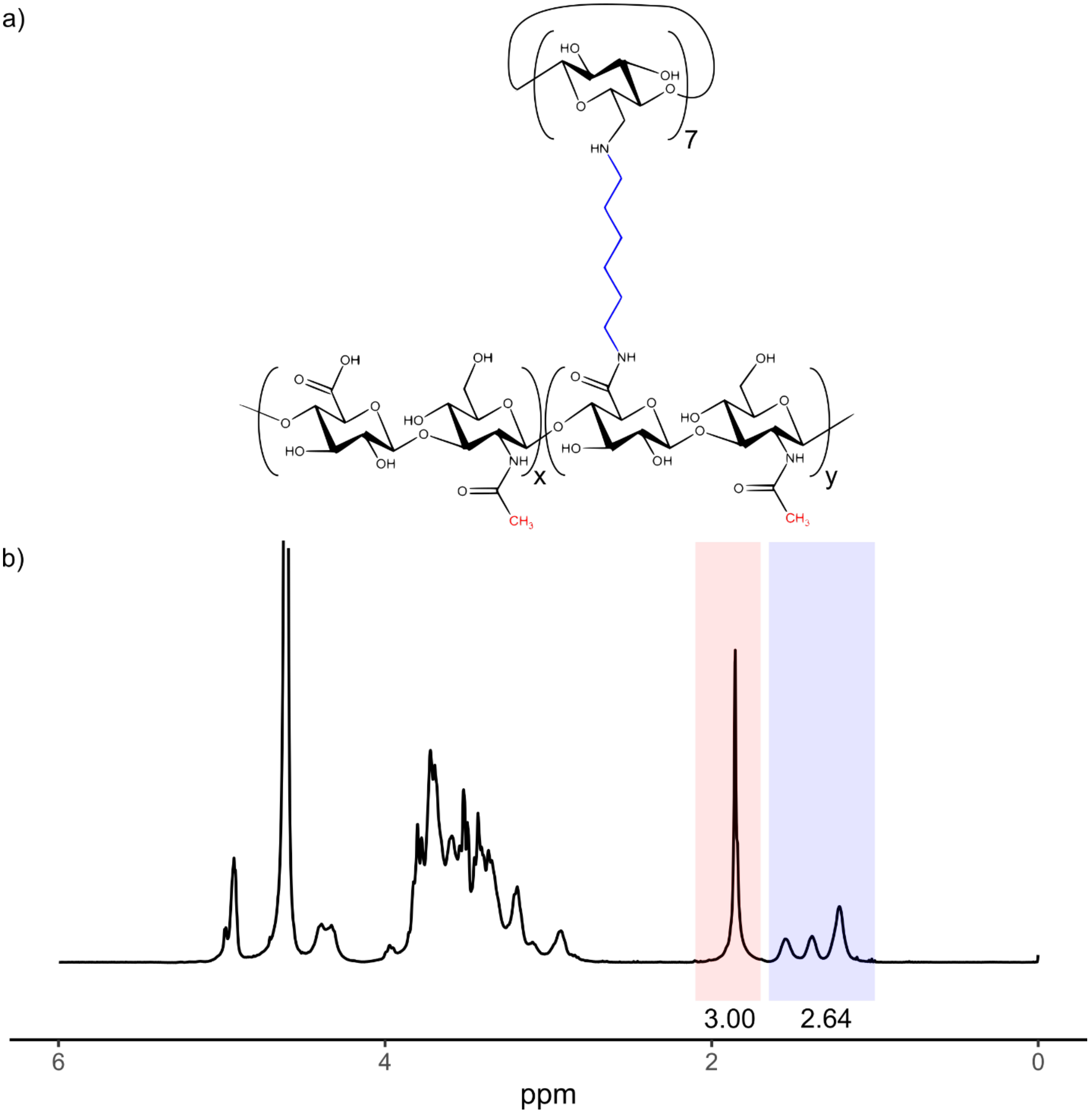


**Figure S1. $^{1}$H NMR spectrum of β-cyclodextrin-modified hyaluronic acid (CDHA).** a) Chemical structure of CDHA with the polymer backbone on lower portion of schematic and pendant cyclodextrin on the upper portion. b) The degree of modification was determined to be 22%. This was calculated from the integration of the hexane linker (δ = 1.00-1.65, 12H) highlighted in blue relative to the methyl group on HA (δ = 1.70-2.10, 3H). Peak areas were normalized such that the HA methyl peak area was equal to 3, and the relative peak area of the hexane linker was divided by 12 to yield the modification.

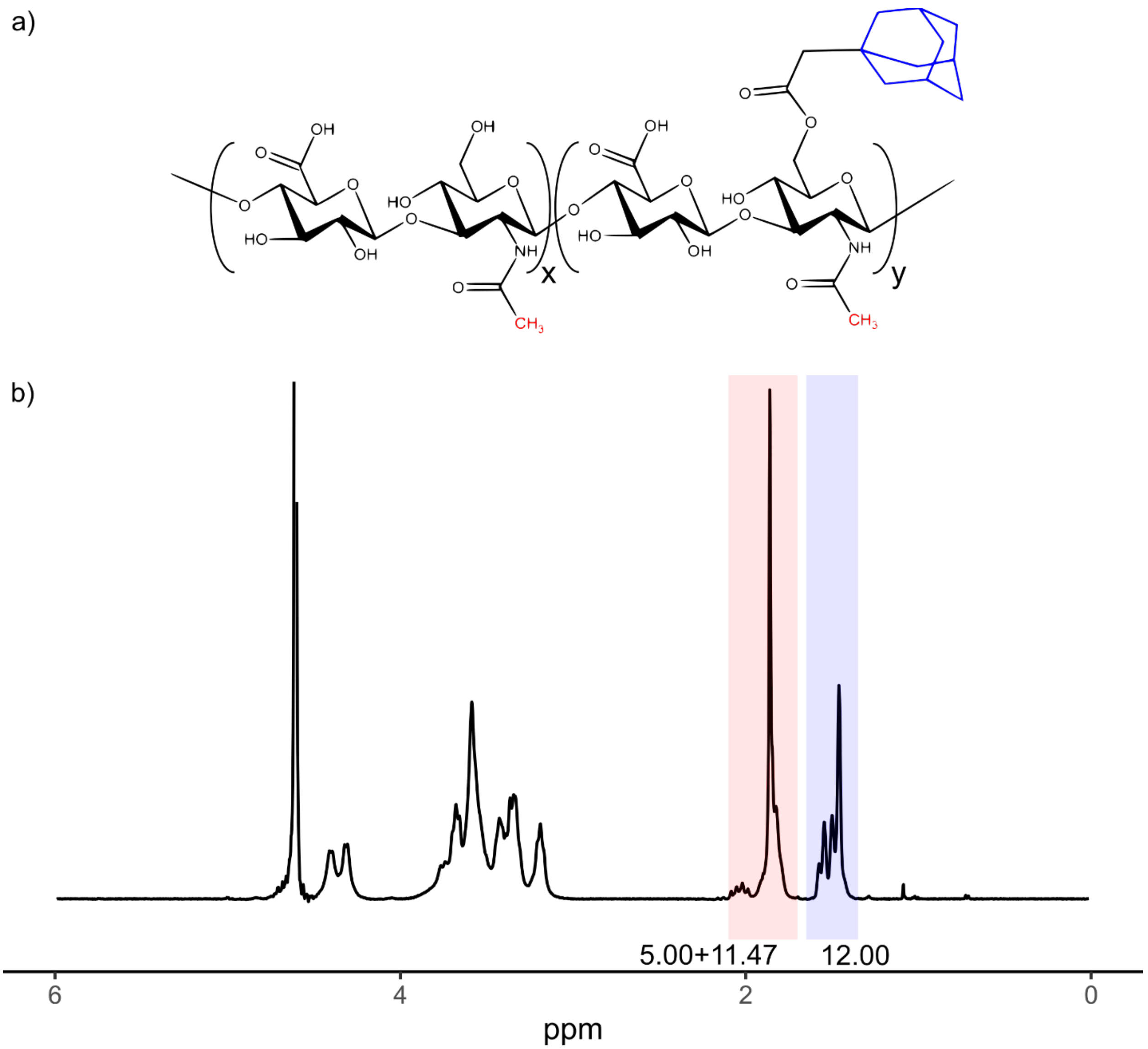


**Figure S2. $^1$H NMR spectrum of adamantane-modified hyaluronic acid (AdHA).** a) Chemical structure of AdHA. b) The degree of modification was determined to be 26%. This was calculated from the integration of the adamantane ethyl groups (δ = 1.35-1.65, 12H) highlighted in blue relative to the peak containing both the methyl group on HA as well as the remaining 5 hydrogens of the adamantaneacetic acid (δ = 1.70-2.10). Peak areas were normalized such that the adamantane ethyl group peak had an area of 12. Then 5 was subtracted from the other normalized peak area, yielding the peak area attributed solely to the methyl group on HA. Peak areas were then renormalized such that the HA methyl peak area was equal to 3, and the relative peak area of the adamantaneacetic acid ethyl groups was divided by 12 to yield the modification (3/11.47*12/12=0.262).

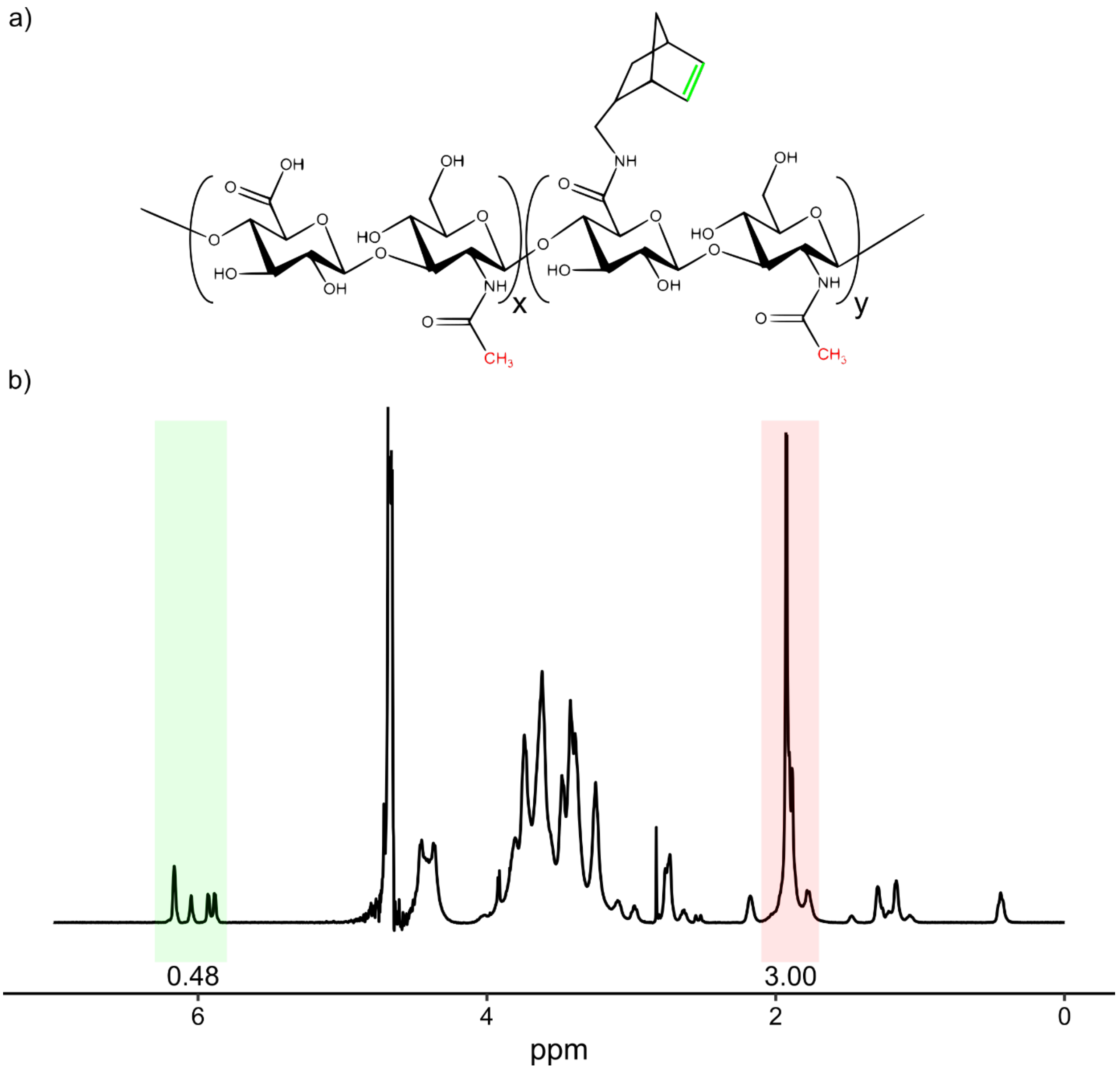


**Figure S3. $^{1}$H NMR spectrum of norbornene-modified hyaluronic acid (NorHA).** a) Chemical structure of NorHA. b) The degree of modification was determined to be 24%. This was calculated from the integration of the norbornene (δ = 5.80-6.25, 2H) highlighted in green relative to the methyl group on HA (δ = 1.70-2.10, 3H) highlighted in red. Peak areas were normalized such that the HA methyl peak area was equal to 3, and the relative peak area of the norbornene group was divided by 2 to yield the modification.

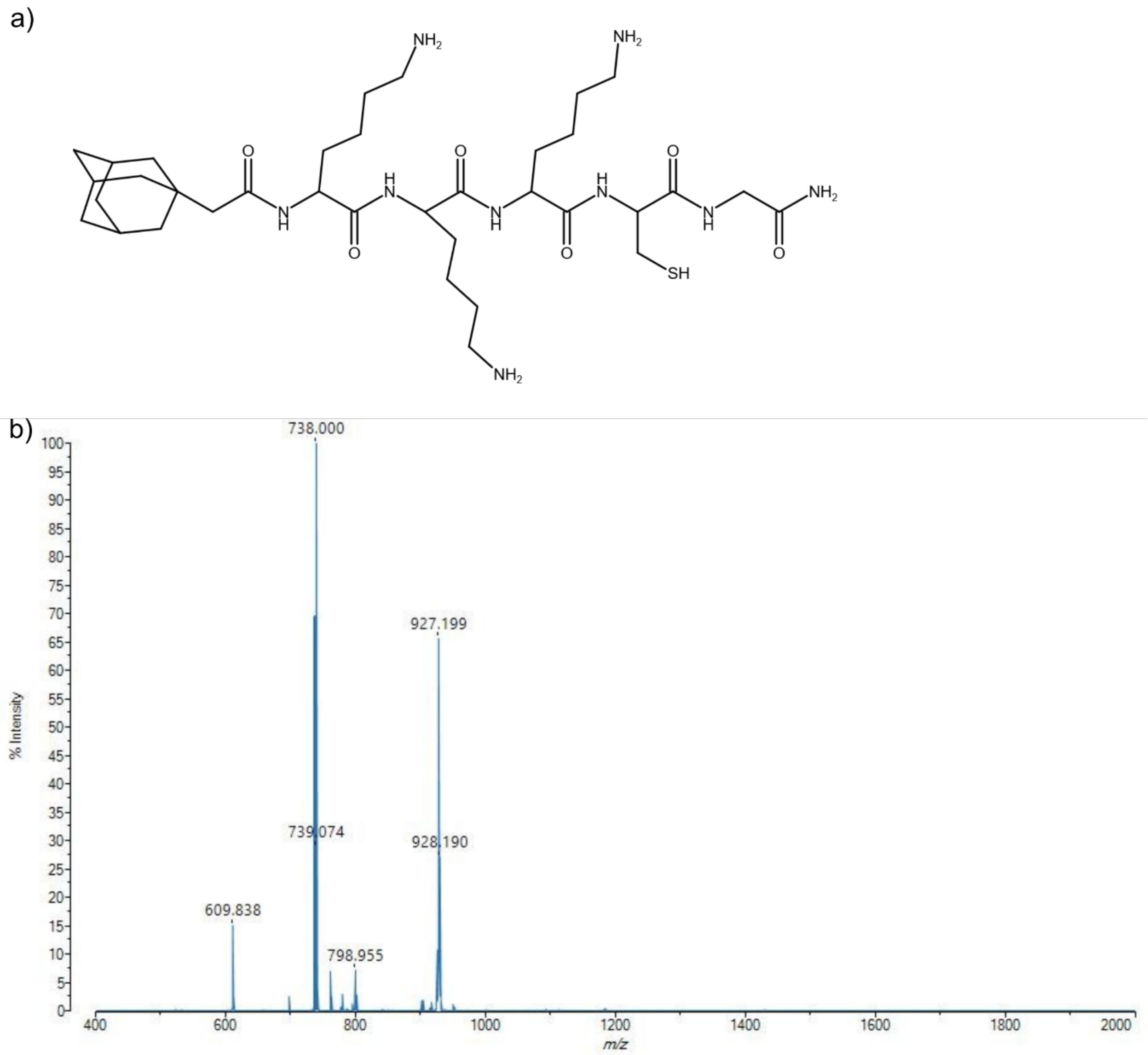


**Figure S4. MALDI spectrum of thiolated adamantane peptide with the sequence 1-adamantaneacetic acid-KKKCG.** a) Chemical structure of synthesized peptide. b) MALDI spectrum of peptide after treatment with TCEP. Expected mass: 738 g/mol. Actual mass: 738 g/mol. The other major product at 927 g/mol is likely an adduct formed between the peptide and the CHCA matrix used, resulting in an ion with a +189 g/mol mass shift[1]. The minor product at 610 g/mol likely indicates a small portion of peptide with a single dropped lysine (-128 g/mol mass shift). No dimerization was detected (expected mass 1474 g/mol).

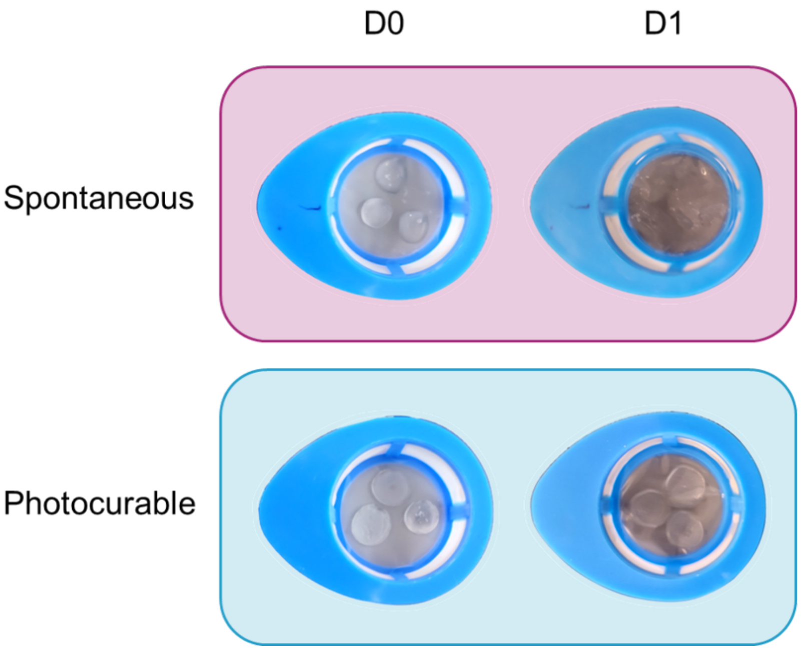


**Figure S5. Hydrogel degradation before and after incubation in excess adamantane.** Hydrogels were incubated in PBS overnight to swell, then transferred to an adamantane solution for another overnight incubation. The photocured hydrogels maintained their shape, while the spontaneously-crosslinked hydrogels disassociated into a single thin layer.

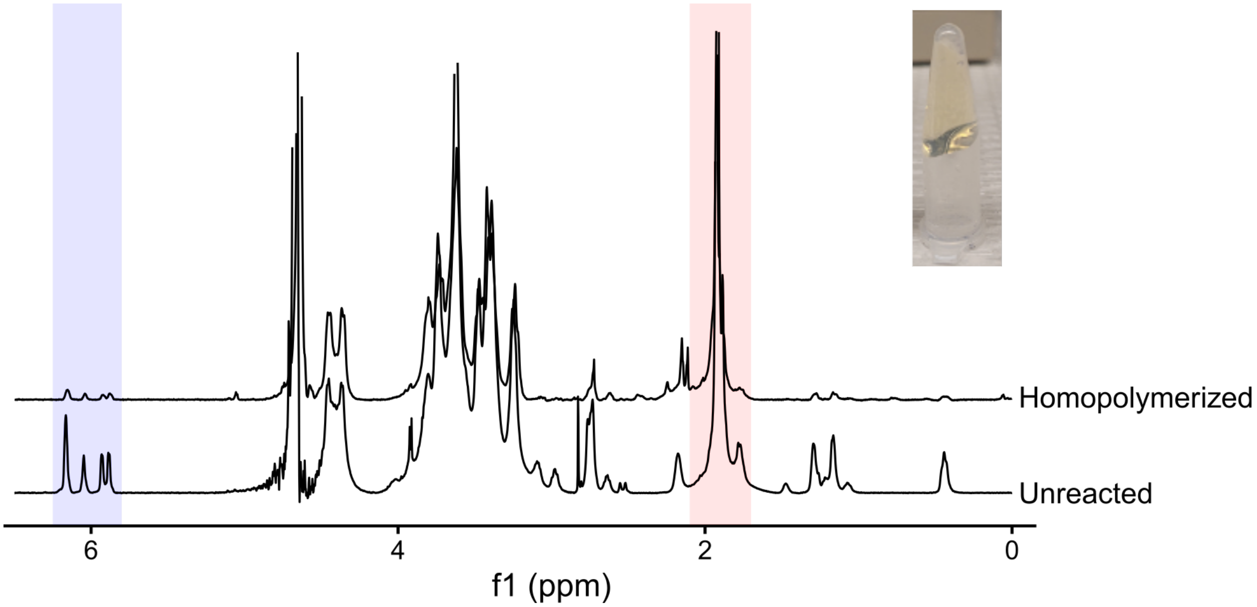


**Figure S6. $^{1}$H NMR spectrum of NorHA before and after exposing to UV light in the presence of LAP photoinitiator.** Remaining norbornene was calculated by integrating the norbornene (δ = 5.80-6.25, 2H) highlighted in blue relative to the methyl group on HA (δ = 1.70-2.10, 3H) highlighted in red. The normalized norbornene peak area of the homopolymerized NorHA (0.104) was divided by the normalized norbornene peak area of unreacted NorHA (0.459), yielding 77% of norbornenes reacted despite absence of thiol. The inset picture depicts the homopolymerized hydrogel prior to digestion by hyaluronidase.

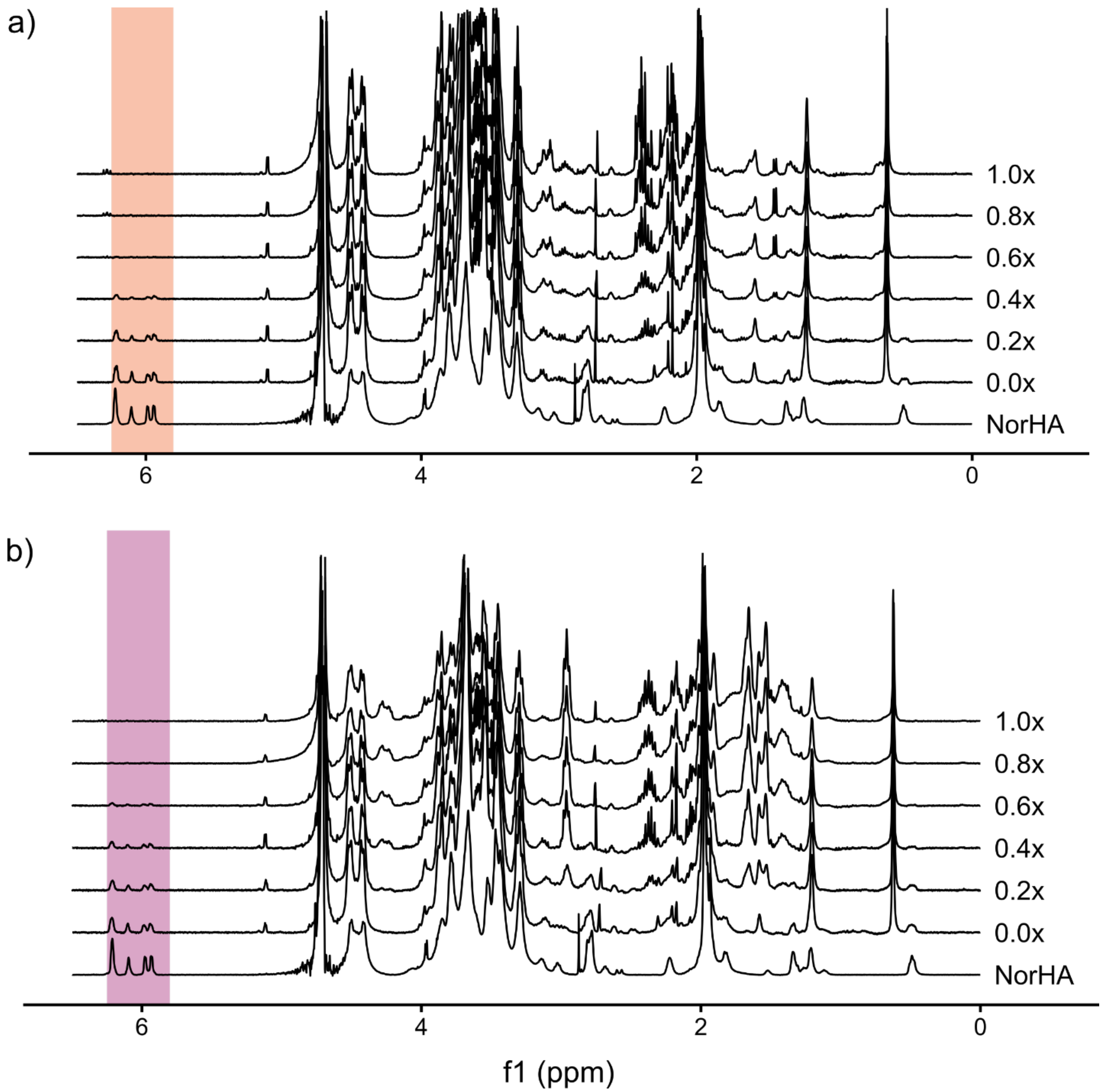


**Figure S7. $^1H$ NMR spectra of NorHA before and after exposure to UV light in the presence of LAP photoinitiator and increasing concentrations of a) cysteine or b) thiolated adamantane.** Norbornene conversion in the presence of thiol (0.2x indicates a 0.2:1.0 ratio of thiol:norbornene). a) Remaining norbornene was calculated by integrating the norbornene (δ = 5.80-6.25, 2H) relative to the methyl group on HA (δ = 1.70-2.10, 3H). These normalized peak areas were divided by the normalized norbornene peak area of unreacted NorHA (bottom spectrum). b) Remaining norbornene was calculated by integrating the norbornene (δ = 5.80-6.25, 2H) relative to the methyl group on HA (δ = 1.70-2.10, 3H). These normalized peak areas were divided by the normalized norbornene peak area of unreacted NorHA (bottom spectrum).

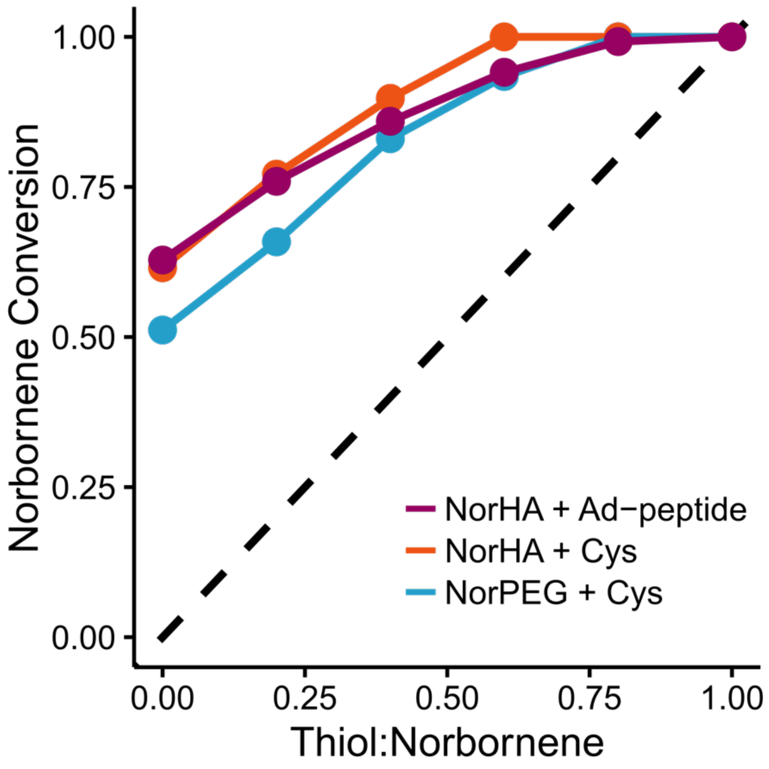


**Figure S8. Proportion of converted norbornene in either NorHA or NorPEG solution with 11.0 mM norbornene, 4 mM LAP, and increasing cysteine or thiolated adamantane concentration.** All conditions exhibited similar norbornene conversion, demonstrating reproducibility of unexpected excess norbornene conversion.

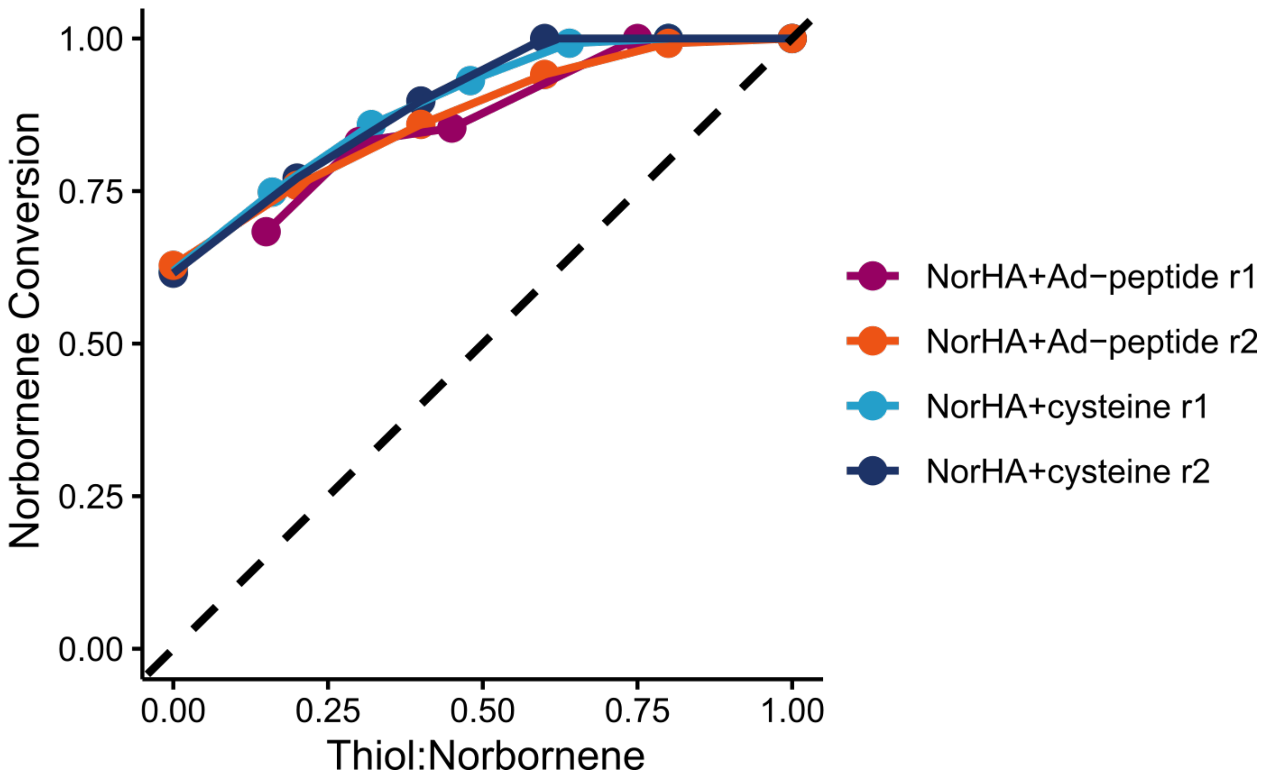


**Figure S9. Proportion of converted norbornene in NorHA solution with 11.0 mM norbornene and increasing thiol concentration.** Replicates within conditions show high overlap. There is a potential difference between the two thiols, though conversion rates are roughly equivalent. The dotted line indicates the expected curve of a 1:1 thiol-ene addition. The samples labeled with "r2" are reproduced from Figure S8.

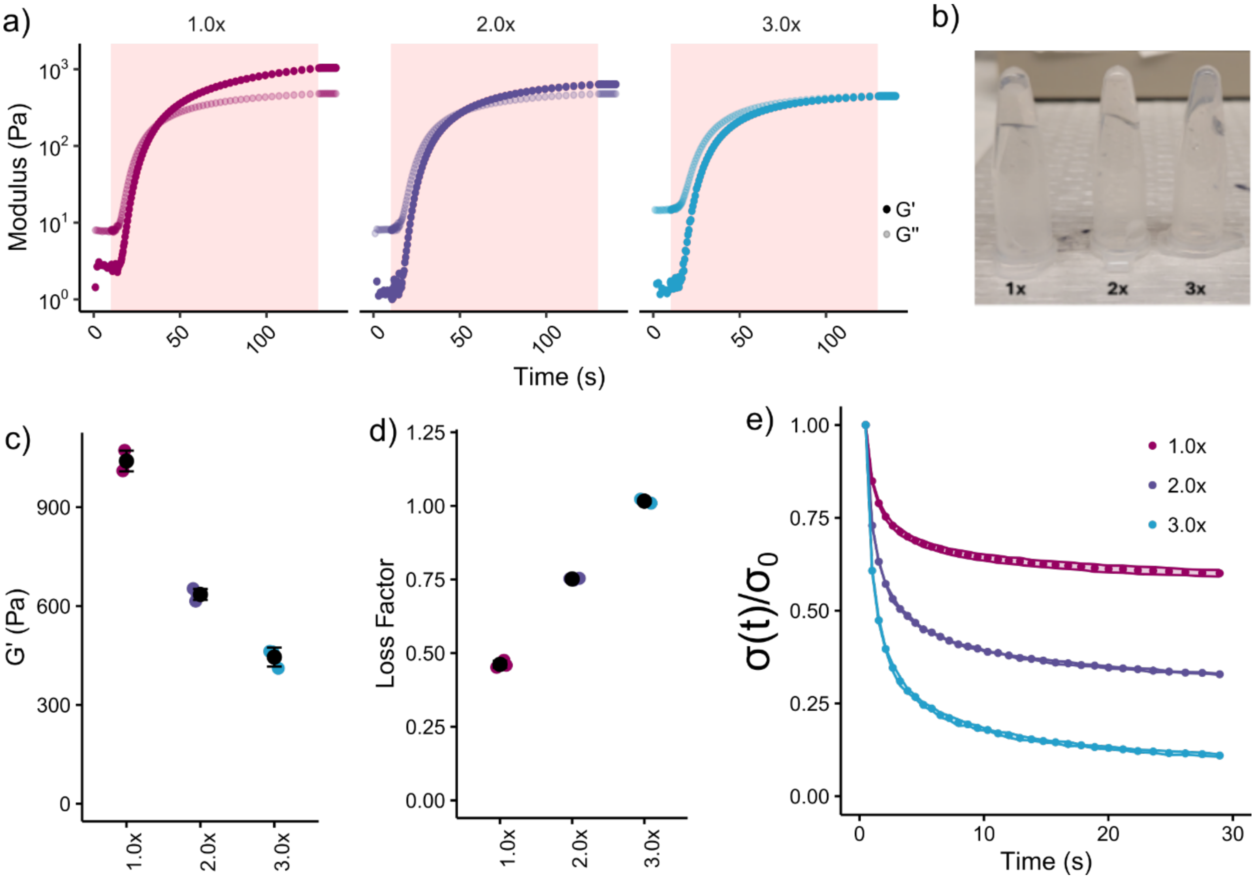


**Figure S10. Rheological characterization of photocurable system with excess adamantane peptide.** a) Monitoring storage (G') and loss (G'') modulus during curing of system with 1:1 thiol:norbornene (1.0x), 2:1 thiol:norbornene (2.0x), and 3:1 thiol:norbornene (3.0x). b) The formulations become increasingly liquid-like, demonstrating a decrease in solid-like behavior as thiol content increases. c) Storage modulus decreases as thiol content increases, while loss modulus is constant, ranging from 450-475 Pa. This suggests a decrease in norbornene homopolymerization. d) Similarly, loss factor increases as thiol content increases. e) Excess thiol enables a greater extent of stress relaxation (stress over time, σ(t), normalized to initial stress, σ(0)), as presence of static covalent bonds from norbornene homopolymerization is reduced.

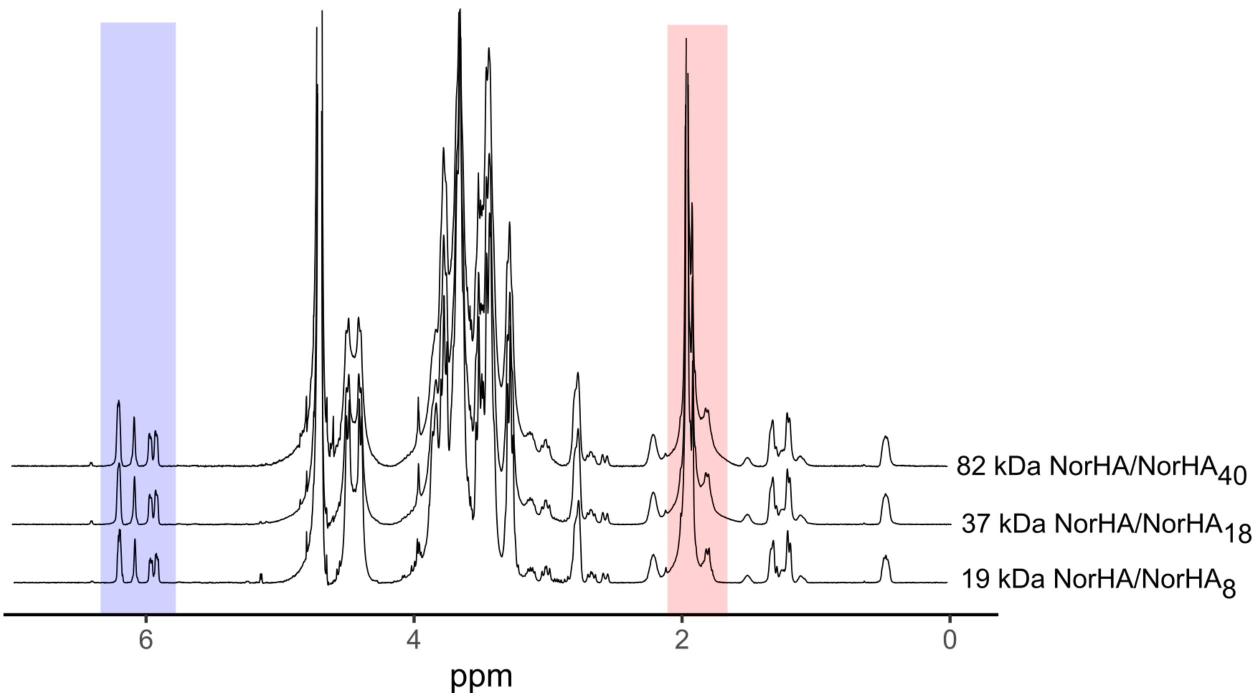


**Figure S11. $^1$H NMR spectra of NorHA synthesized from different molecular weights of HA.** The degree of modification was determined to be 17% for 19 kDa, 18% for 37 kDa, and 18.5% for 82 kDa. This was calculated from the integration of the norbornene (δ = 5.80-6.25, 2H) highlighted in blue relative to the methyl group on HA (δ = 1.70-2.10, 3H) highlighted in red for each polymer.

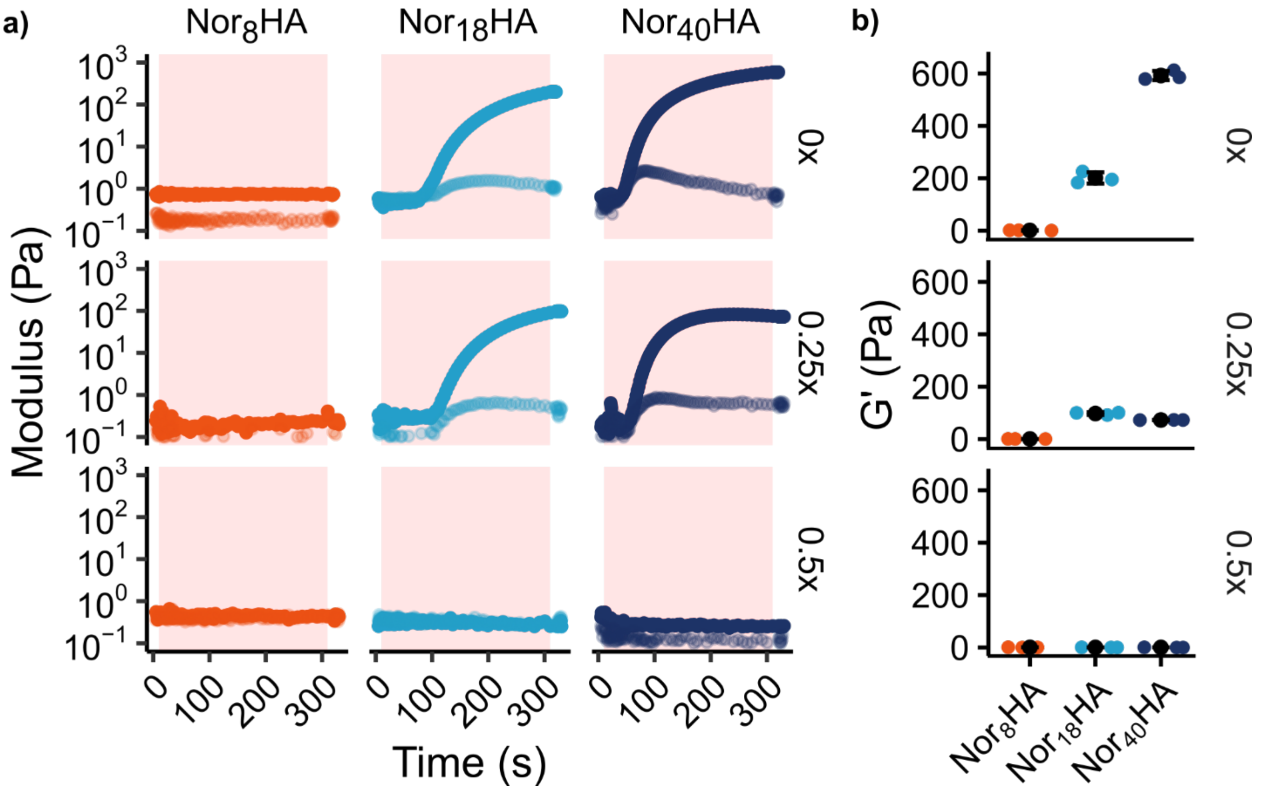


**Figure S12. Reducing f while maintaining norobornene concentration inhibits network formation by norbornene homopolymerization.** a) Oscillatory rheological test monitoring UV curing of different NorHA polymers (columns) with different molar ratios of cysteine (rows) and 4 mM LAP photoinitiator. $Nor_8HA$ did not gel even at 0 mM thiol concentration, while increasing thiol reduced gelation capability of the other two polymers. b) Final storage modulus after 5 minutes of UV exposure.

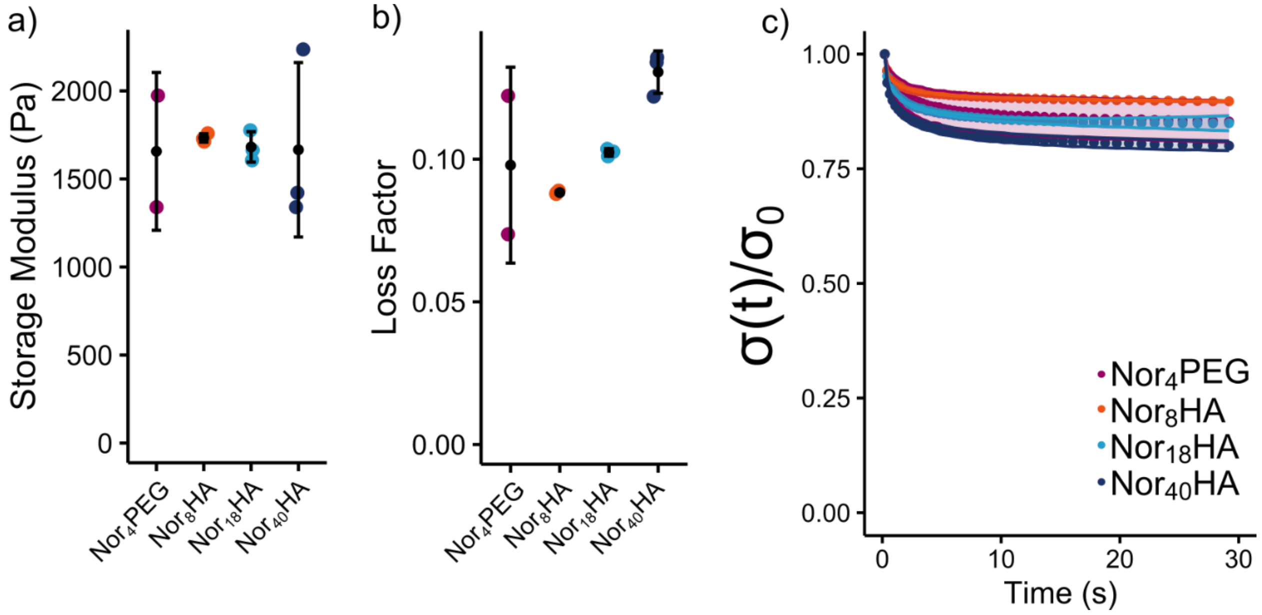


**Figure S13.** ***In situ*** **rheological characterization of hydrogels with similar mechanics but different polymers.** a) Storage moduli after 5 minutes of UV exposure show similar mean G' regardless of polymer. b) Loss factor after 5 minutes of UV exposure increases slightly as f increased. c) Stress relaxation tests showed relaxation extents ranging from 10%-20%, with higher f polymers relaxing slightly more.

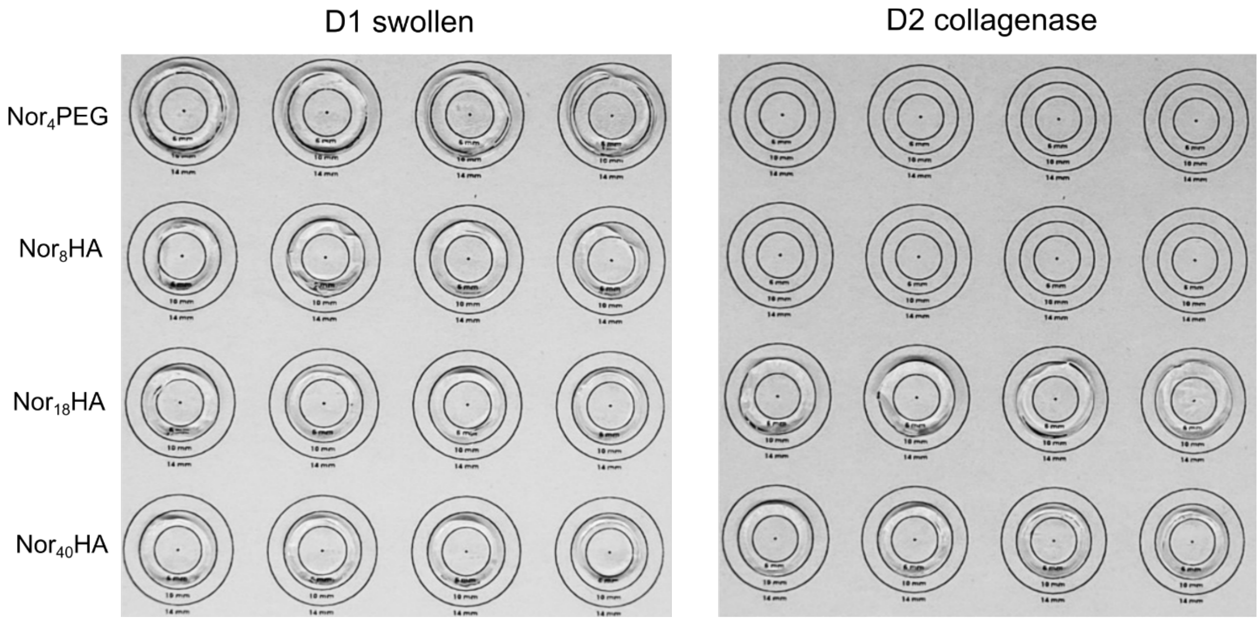


**Figure S14. Degradation of hydrogels of similar mechanics by collagenase.** Hydrogels were incubated in PBS overnight to swell, then transferred to a collagenase solution (150 U/mL) for another overnight incubation. The $NorHA_{40}$ gels did not contain any MMP-degradable crosslinker, and thus did not degrade. The $NorHA_{18}$ hydrogels were still intact despite using a high concentration of collagenase which should have cleaved all crosslinker. The $Nor_4PEG$ and $NorHA_8$ hydrogels were completely disassociated.
